# Organelle communication networks rewire to support lipid metabolism during neuronal differentiation

**DOI:** 10.64898/2026.02.10.704675

**Authors:** Maria Clara Zanellati, Zachary Coman, Disha Bhowmik, Chih-Hsuan Hsu, Richa Basundra, Shannon N. Rhoads, Ngudiankama R. Mfulama, Brandie M. Ehrmann, Mohanish Deshmukh, Sarah Cohen

## Abstract

Cell fate transitions require coordinated remodeling of intracellular organelles, but how organelle morphology and interactions rewire during neurogenesis remains unclear. Here we combine multispectral imaging with quantitative organelle signature analysis to simultaneously map eight organelles as human induced pluripotent stem cells differentiate into forebrain-like neurons. We find compartment and time-specific rescaling of organelles and a progressive increase in higher-order membrane contacts, with mitochondria emerging as an early interaction hub. Later, endoplasmic reticulum (ER)-organelle contacts dominate with ER-peroxisome contacts promoting ether lipid biosynthesis, membrane homeostasis and synapse formation. Disrupting this contact impairs plasmalogen production, synaptic organization, and neuronal activity, identifying the ER-peroxisome axis as a key regulator of neuronal maturation.

## Main Text

Cell fate transitions are accompanied by extensive remodeling of intracellular organelles to support specialized physiological functions [1, 2]. These include changes in organelle size, abundance, morphology and distribution. Neuronal differentiation is an example of a cell fate change that requires dramatic changes in cell morphology and function. Transcriptomic, proteomic and microscopy studies have shown that neurogenesis is associated with a switch from glycolytic to oxidative metabolism and large-scale reorganization of the mitochondria, endoplasmic reticulum (ER), Golgi, and endolysosomal compartments, driven in part by selective autophagy [3–9]. However, previous studies focused on individual organelles or pathways, leaving unresolved how organelles as a coordinated system rewire to support differentiation of pluripotent stem cells into neurons.

Recent work discovered that in addition to having different organelle landscapes, the pattern of interactions among organelles is distinct for specialized cell types such as neurons and astrocytes [10]. Organelles interact at membrane contacts sites (MCSs): sites of close apposition mediated by protein tethers where organelles including ER, mitochondria, Golgi, endolysosomes, peroxisomes and lipid droplets exchange metabolites [11–13]. Multispectral imaging (MSI) has emerged as a powerful approach to resolve organelle organization and interactions, enabling robust discrimination of cell states under basal conditions and in response to stress [10, 14–16]. We reasoned that organelle interaction networks must emerge during differentiation and that mapping the organelle interactome across differentiation would offer the opportunity to uncover previously unappreciated layers of organelle regulation.

Here, we use MSI together with quantitative image analysis and lipidomics to follow how eight organelles remodel as human induced pluripotent stem cells (iPSCs) differentiate into forebrain-like neurons (iNeurons). We find that differentiation is accompanied by a global rescaling of organelles, with distinct geometries in soma versus neurites. We further reveal a progressive increase in organelle connectivity, with higher-order (3- and 4-way) contacts forming as neurons mature. We identified a sequential reorganization of the mitochondria and lipid-related interactomes at distinct stages of differentiation. Finally, we show that ER-peroxisome contacts promote plasmalogen biosynthesis and are required for proper synapse formation and neuronal activity.

### Multispectral imaging reveals temporal organelle remodeling during neuronal differentiation

We generated human iPSCs [17] stably integrated with the transcription factor hNGN2 that efficiently differentiate into forebrain-like neurons in less than 14 days (**Fig. S1A-F**). We utilized multispectral imaging, as previously described [10, 14, 15] but with the addition of markers for nuclei and plasma membrane, as well as transmitted light, to quantify morphological changes in eight organelles during neuronal differentiation (**Fig. 1A**; **Fig. S2A-C**). To determine whether expression of tagged constructs perturbed organelle homeostasis, we performed Airyscan imaging on fixed samples stained for endogenous markers for each organelle (excluding Golgi) at matched time points (**Fig. S3A-N**). We did not observe morphological distortions, indicating that our labeling strategy does not grossly perturb organelle organization under these conditions.

**Figure 1:**
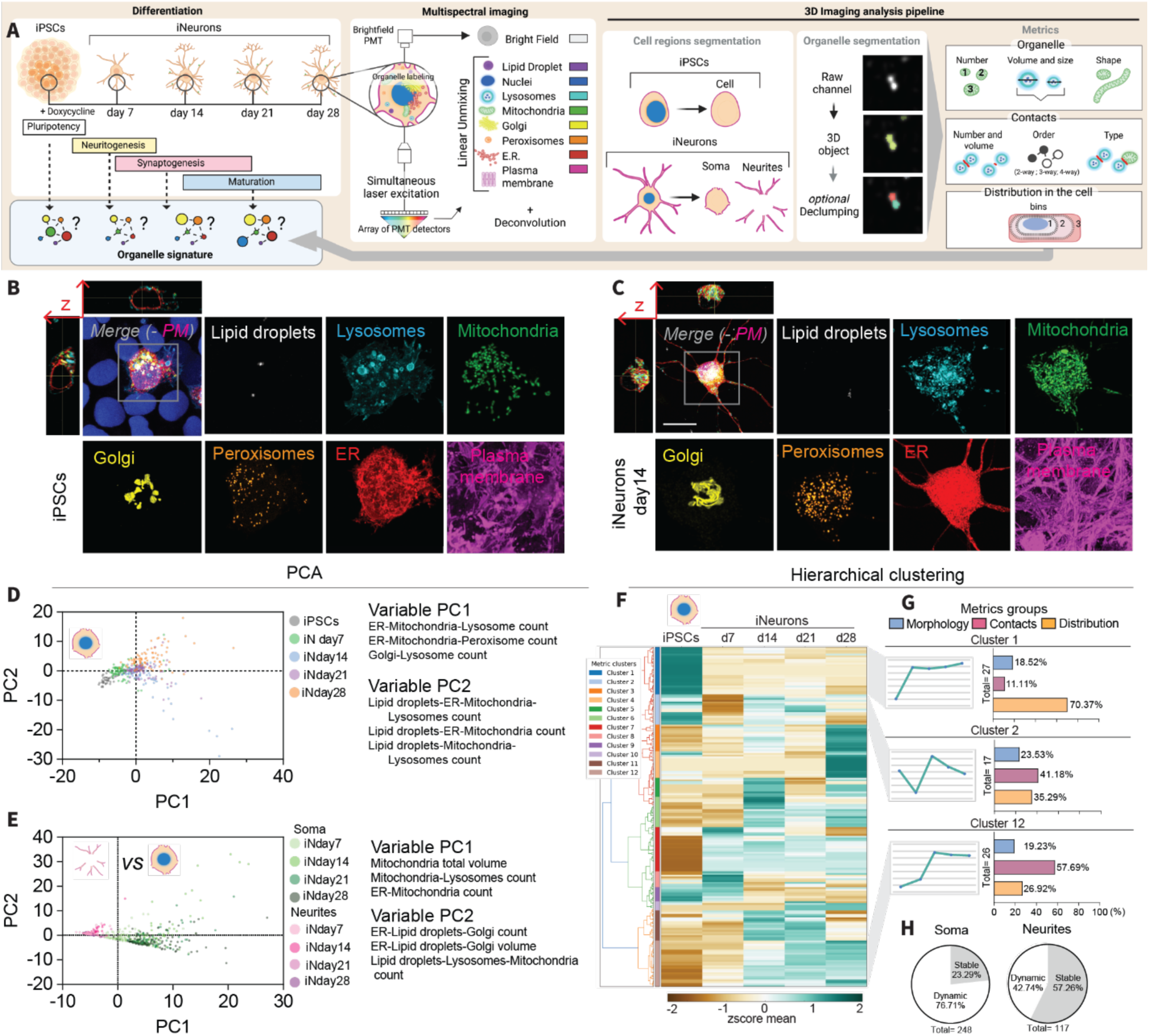
Multispectral imaging as a tool to reveal organelle rewiring during differentiation. (**A**) Schematic representation of experimental design, using rapid differentiation into forebrain like neurons. (**B**, **C**) Deconvolved multispectral images (maximum intensity projection) of iPSCs and iNeurons at day14, showing orthogonal projection (in XY and ZY) without nuclei (blue) and plasma membrane channels, together with individual channels showing different organelles: lysosomes (cyan), mitochondria (green), lipid droplets (white), Golgi (yellow), peroxisomes (orange), ER (red) and plasma membrane (magenta). Scale: 10 µm. (**E**) PCA plots of the first pass metrics (n=248) of iPSCs and iNeurons at different time points (day7, day14, day21, day28). (**D**) PCA plots of the first pass metrics (n=114) of iNeurons and their neurites. (**F**) Hierarchical clustering of 191 out of 248 metrics of iPSCs and iNeurons (soma region). (**G**) Curves representing the dynamics of a representative metric and graphs showing percentage of each metric group (morphology, contacts and distribution of the organelle) for three featured clusters. (**H**) Graphs showing percentage of metric groups that showed dynamic vs. stable patterns, in the soma regions (with iPSCs) or neurites (*left and right panel, respectively*).

To compare organelle remodeling across differentiation stages in iNeurons, we first segmented all the organelles, then separated soma and neurite compartments and analyzed eight organelles using a custom Python-based image analysis pipeline (inferSubC v2.0): nuclei, lipid droplets lysosomes, mitochondria, Golgi, peroxisomes, ER, and plasma membrane (**Fig. 1A-C**; **Fig. S2B, C**). In total, we extracted 5,441 single-cell metrics, including organelle volume, number, and shape, pairwise and multiple organelle contacts, and spatial distribution of organelles and their contacts. To reduce redundancy, we focused subsequent analyses on a curated subset of these metrics for statistical comparison (**Data S1-3**). Principal component analysis (PCA) of soma metrics revealed clear separation of all differentiation time points, with the major components dominated by contact metrics (**Fig. 1D**). In contrast, PCA comparing soma and neurites showed a robust separation between these two compartments irrespective of time point, indicating that soma and neurites represent distinct organelle states (**Fig. 1E**). When analyzed alone, neurites showed limited separation by PCA (**Fig. S4A**), suggesting that the most pronounced remodeling occurs in the cell body.

We next examined the temporal dynamics of these metrics by hierarchical clustering (**Fig. 1F-H**; **Fig. S4B-E; Data S4**, **5**). Metrics segregated into 12 clusters with distinct trajectories across differentiation. The largest clusters, e.g., clusters 1 and 7 (**Fig. 1F, G, Fig. S4Cv**), showed pronounced changes during the transition from iPSCs to neurons, followed by relative stabilization, and were enriched for distribution-related metrics (**Fig. 1F, G**), consistent with early establishment of cellular polarity (**Fig. S5A-Q**). Notably, most organelles progressively redistributed toward the basal region of the cell as iPSCs differentiated into iNeurons (**Fig. S5C, E, G, I, K, P**), whereas lipid droplets displayed the opposite trend (**Fig. S5M, Q**), potentially reflecting a shift from lipid storage toward lipid mobilization for membrane expansion or secretion. In parallel, several organelles shifted toward the cell periphery, consistent with increased trafficking into developing neurites (**Fig. S5D, F, H, J, L, N**, and **P**, **Q**). A second type of cluster, e.g., cluster 12, was enriched for contact-related metrics, increased from day 7 to day 14 and remained elevated through day 28. By contrast, cluster 2, enriched for morphology and contact metrics, decreased at day 7, peaked at day 14, and declined thereafter. Together, these dynamic patterns suggest a sequential and temporally coordinated reorganization of organelle architecture during neuronal differentiation.

Whereas soma metrics displayed rich and dynamic temporal patterns, neurite metrics were comparatively homogeneous, with only a subset of features transiently peaking at day 14 before decreasing by day 28 (**Fig. 1H**; **Fig. S5D-E**). Together, these data reveal extensive, temporally structured organelle remodeling during neuronal differentiation, which is more pronounced in the cell body than in neurites.

### Organelles rescale during differentiation

As iPSCs differentiated into iNeurons, we observed an initial reduction in soma volume at day 7 (**Fig. 2A**), accompanied by an increase in both the number and total volume of neurites (**Fig. 2B, C**). Across this process, most organelles scaled their total volumes proportionally with soma or neurite volume, indicating a global rescaling of organelle content with cell geometry (**Fig. 2D-J**; **Fig. S6A,B, J-P**). A notable example was the ER, whose total volume significantly contracted at day 7 followed by an expansion at day 14 (**Fig. 2E**). This is consistent with enhanced ER-phagy during early differentiation, as reported previously [4, 5], and is further supported by an increase in ER tubules (**Fig. S6R**). A similar trend was observed for the nucleus, lysosomes, Golgi and peroxisomes (**Fig. 2G-I**). Peroxisome total volumes rose as iNeurons matured, due to an increase in their number/cell (**Fig. S6N**), despite reducing in individual size (**Fig. 2N**; **Fig. S6T**), as previously observed during neuronal differentiation and development [18–20]. Although not significant, lipid droplets increased their total mass reaching a peak at day 14 and then decreasing, perhaps suggesting a supportive role in fatty acids and cholesterol for membrane growth [21], in line with the distribution data (**Fig. 2J**; **Fig. S6N, O, and Q**). In contrast, mitochondrial volume and number increased during differentiation, especially in the soma region (**Fig. 2F**; **Fig. S6B, F**), together with an increase in complexity as shown from an increase in surface area to volume ratio (**Fig. S6S**), consistent with a shift toward oxidative phosphorylation and increased mitochondrial reliance in neurons [7–9, 22].

**Figure 2:**
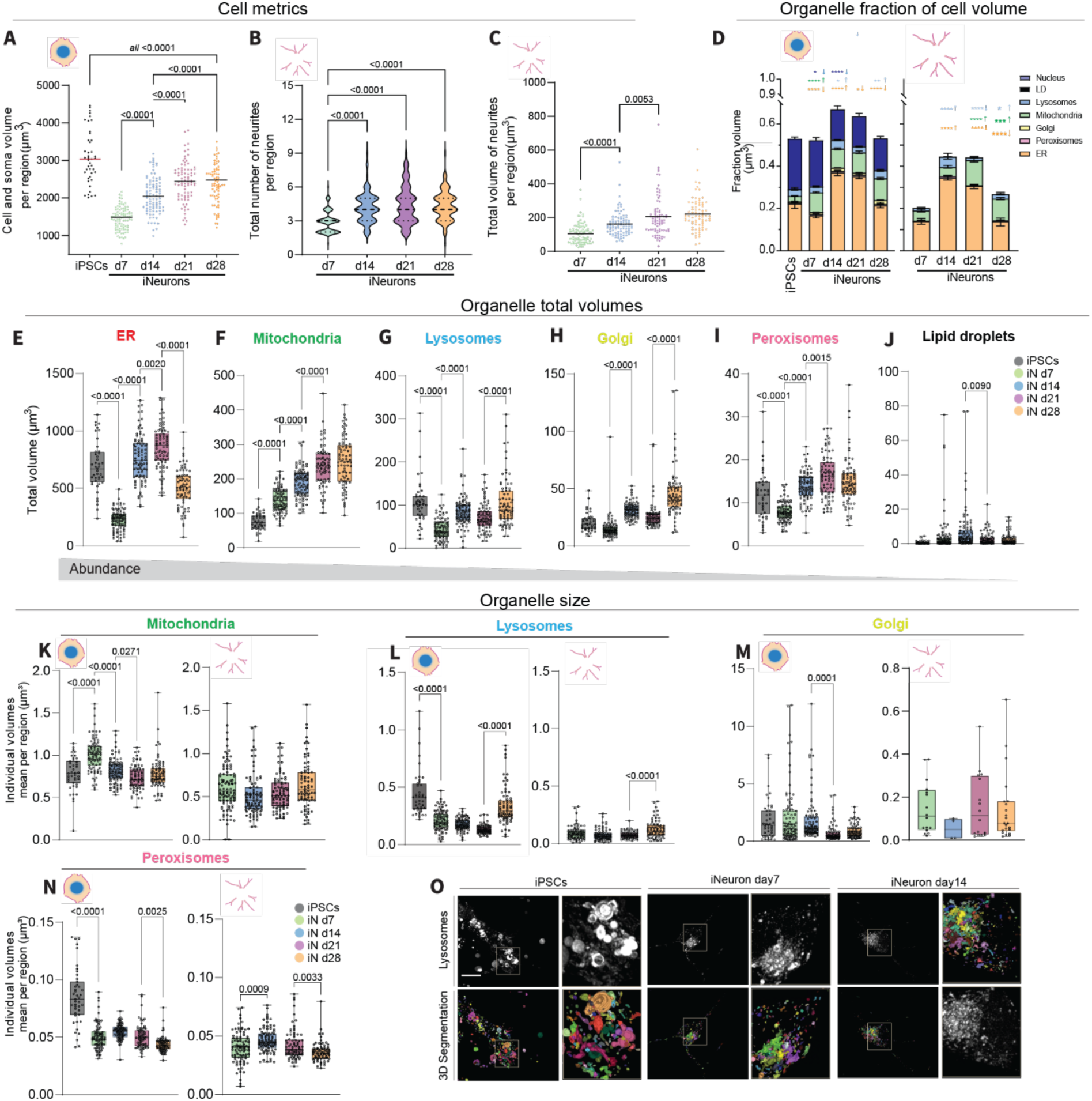
Dynamic temporal and spatial rescaling of organelles during neuronal differentiation. (**A**) iPSC whole cell and iNeuron soma region total volumes (µm^3^) per cell. (**B**) Total volumes and (**C**) number of neurite regions in iNeurons at different time points (day7, day14, day21, and day28). (**D**) Organelle total volumes in iPSCs and iNeurons normalized to soma or neurite volumes (*left* and *right panel*, respectively). (**E**-**J**) Organelle total volumes of iPSC and iNeuron soma regions. (**K**-**N**) Individual organelle size in iPSCs and iNeurons (soma and neurites). (**O**) Deconvolved images and 3D segmentations of the lysosome channel in iPSCs and iNeurons at day7 and day14. Scale: 10 µm. Two-way ANOVA, p-value versus previous time point.

For organelles with highly variable size in iPSCs, such as lysosomes, we observed a decrease in individual organelle volume, likely facilitating their transport into thin neuronal processes (**Fig. 2L, O**; **Fig. S6D, E, U**). By contrast, organelles with a more stereotyped size, such as peroxisomes and lipid droplets, displayed similar individual volumes in soma and neurites (**Fig. 2C**). Moreover, Golgi differed in size between soma and neurites, consistent with the existence of compartment-specific organelle populations (**Fig. 2M**) [6, 23]. In many cases, a reduction in individual organelle size was accompanied by an increase in organelle number (**Fig. S6E-I, Q**). These results were further validated by AiryScan imaging, with the exception of mitochondria organelle number (**Fig. S3B-N**). Together, these data show that organelles rescale and remodel in a compartment and time-specific manner during neuronal differentiation, establishing a new organelle landscape that likely underpins the emergence of neuron-specific functions and interaction networks.

### Inter-organelle interactions increase as neurons differentiate

We reasoned that as organelles remodel, so would their inter-organelle communication networks. Given the pronounced changes in organelle volume and number, we first checked what fraction of each organelle population engages in contacts versus remaining isolated (**Fig. S7A-C**). Across all time points, the majority of organelles were engaged in at least one contact, with a smaller fraction remaining contact-free, in both soma and neurites (**Fig. S7B, C; Data S6**). Moreover, contacts involving lysosomes, mitochondria, Golgi and peroxisomes increased as iPSCs differentiated into iNeurons, whereas lipid droplet contacts decreased in the soma but showed a modest, non-significant increase in neurites. Interestingly, organelles such as lipid droplets and lysosomes consistently showed a lower propensity to engage in contacts across differentiation, indicating organelle-specific interaction frequencies that are largely independent of developmental stage. Overall, the likelihood that an organelle engaged in contacts was not strictly determined by its size, with few exceptions (**Fig. S7D, E; Data S6**).

We next asked whether any organelle assumes a more central role at specific stages of differentiation. To assess this, we quantified how often each organelle type contacted each of the others at least once per cell (“organelle partner prevalence”; **Fig. S7A; Data S6**). Independent of time point, each organelle type contacted all other organelle types, but with different magnitudes, and the percentage of organelles engaged in contacts initially increased as iPSCs differentiated into iNeurons in both soma and neurites (**Fig. S7F, G**). This trend suggested a rise in simultaneous interactions between individual organelle objects during differentiation. To test this directly, we measured the “organelle contact degree”, defined as the number of distinct organelle types that a given organelle object contacts at least once, from degree 1 (one type) up to degree 5 (five types) (**Fig. S7A; Fig. 3A; Data S6**). As expected, we observed a progressive increase in the number of organelle objects with higher contact degree across differentiation. Similar to the organelle partner prevalence, the contact degree analysis revealed an early increase in higher-degree interactions among lipid droplets, lysosomes, mitochondria, and peroxisome as iPSCs transition to iNeurons, followed by a later increase in higher-degree Golgi and ER interactions during iNeurons maturation (**Fig. 3A**).

**Figure 3:**
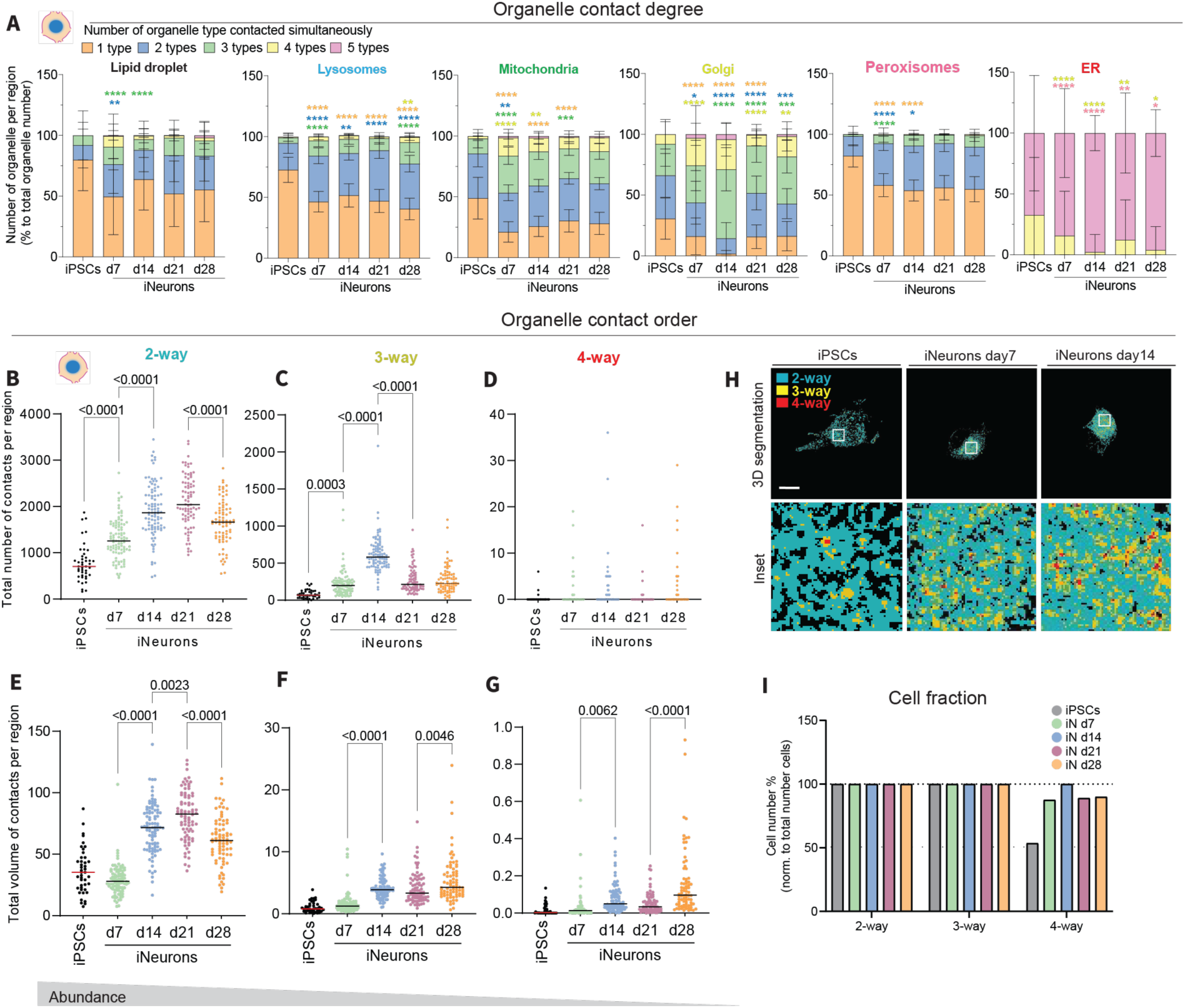
High-order contacts increase as iPSCs differentiate to iNeurons. (**A**) Percentage of the number of organelles with contact degree from 1 to 5, per cell (iPSCs) or soma (iNeurons), for each individual organelle, during neuronal differentiation. (**B**-**D**) Total number or (**E**-**G**) total volume of 2-, 3-, and 4-way contacts per cell in iPSCs and soma region of iNeurons at different time points (day7, day14, day21, and day28). (**H**) 3D segmentations of iPSCs and iNeurons at day14, of all the exclusive 2-way contacts (cyan), 3-way contacts (yellow), and 4-way contacts (red). Scale: 10 µm. (**I**) Percentage of cells that have at least one contact per order (2-,3-, and 4-way), in iPSCs or soma region of iNeurons, at different time points. Two-way ANOVA, p-value versus previous time point.

To further dissect the magnitude and architecture of these interactions, we quantified both the number and volume of contacts between organelle pairs (“contact type”) and classified each contact as involving two, three or, more rarely, four organelles simultaneously, (“contact order”; **Fig. S7A; Data S7**). We observed an overall increase in the total number and volume of contacts per cell, in both soma (**Fig. S8A, B**) and neurites (**Fig. S8C, D**), with contact number rising early as iPSCs differentiate into iNeurons, followed by a subsequent increase in contact volume. This initial increase in contact number, followed by a rise in contact volume, indicates that the organelle network is first established and then strengthened over time. This increase was largely driven by a rise in 2- and 3-way, and more rarely 4-way, contacts during differentiation (**Fig. 3B-H**). While 2-way contacts remained the most abundant, we detected a significant increase in the fraction of 3-way contacts in the soma as iPSCs differentiated into iNeurons, indicating enhanced connectivity within the cell body (**Fig. S8E**). In contrast, neurites showed no detectable change in the contact fractions, consistent with a more homogeneous interaction landscape in this compartment, with the exception of an increase in 2-way contacts when iNeurons reach day 28 (**Fig. S8F-N**). In iPSCs, 4-way contacts were present in only ∼50% of cells, whereas a substantially larger fraction of iNeurons exhibited at least one 4-way contact per cell (**Fig. 3I**), suggesting that the emergence of a more complex, higher-order organelle interactome is a hallmark of neuronal differentiation.

### Specific inter-organelle interactions increase during different stages of differentiation

Many three-way organelle contacts have recently been implicated in organelle biogenesis and cell homeostasis [24–28]. The cellular function of a given tripartite contact is determined by the specific organelle combination involved. We therefore quantified the total number and volume of three-way contacts during differentiation.

We observed ER-mitochondria-lysosome contacts to be the most abundant tripartite assemblies, particularly at early stages of differentiation, followed by ER-mitochondria-peroxisome contacts, in both soma and neurites (**Fig. 4A-C**; **Fig. S9A**). Most of the tripartite contacts that significantly increase as iPSCs differentiate to iNeurons involve mitochondria, again consistent with the central role of mitochondria early during differentiation, and the initial need of metabolic rewiring. To better understand the dynamics underlying these tripartite contacts, we examined pairwise contact number and volume for each organelle pair (**Fig. 4D**; **Fig. S9B**). Given the striking changes in cell and soma volumes we normalized the contact volumes to cell or region volume (**Fig. 4D**). As iPSCs differentiated into iNeurons, we observed a marked increase at day 7 in ER-mitochondria, lysosome-mitochondria, Golgi-mitochondria, and mitochondria-peroxisome contacts in the soma region (**Fig. 4D**). ER-mitochondria and lysosome-mitochondria contacts facilitate mitochondrial fission while the ER is also present at sites of mitochondrial fusion [27, 29–31]. Similarly, Golgi-derived vesicles have been reported to support late steps of mitochondrial fission [32]. Stable Golgi-mitochondria contacts have also been linked to the regulation of calcium gradients, in other cell types [33], but their roles remain largely unexplored. Mitochondria-peroxisome contacts help mitochondria offload damaging reactive oxygen species [34], a function that is likely crucial to sustain the increased mitochondrial activity in iNeurons. Together, these findings suggest that an early and major step in differentiation is a broad remodeling of the mitochondrial interactome, potentially to support a new metabolic state reliant on oxidative phosphorylation.

**Figure 4:**
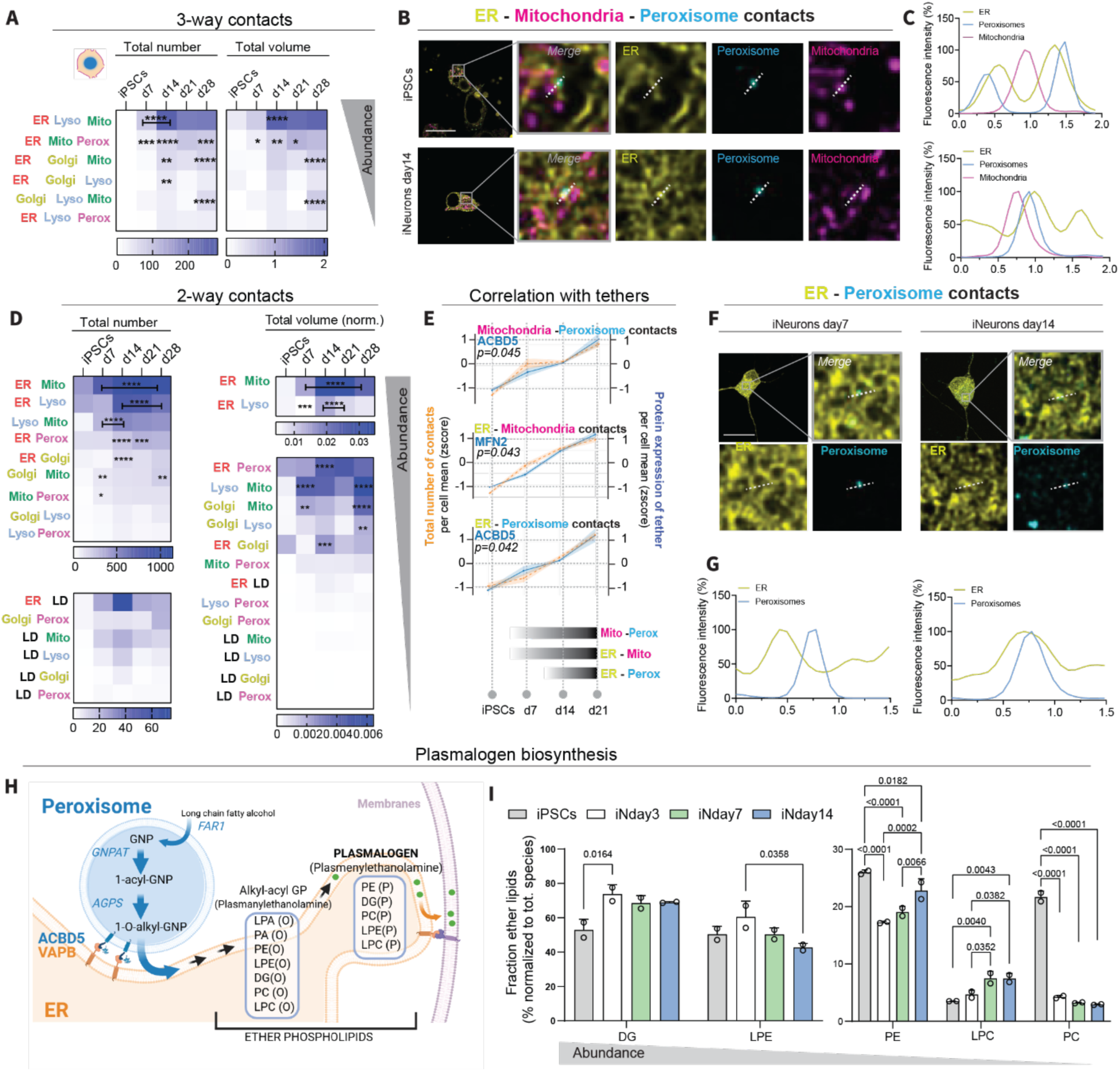
Pairwise and tripartite contacts are sequentially assembled along differentiation. (**A**) Total number (median) of all the 3-way contact types per cell (iPSCs) or soma region (iNeurons) at different time points (day 7, 14, 21 and 28). Two-way Anova; p-value versus previous time point. (**B**) Multispectral images and (**C**) the corresponding line scan quantification, of the tripartite contact between ER, mitochondria and peroxisome, in iPSCs and iNeurons at day 14. (**D**) Total number (median) of all the 2-way contact types per cell (iPSCs) or soma region (iNeurons) at different time points. Two-way ANOVA, p-values versus previous time point. (**E**) Z score of the total number of indicated contacts, together with the expression level of tethering proteins ACBD5 and MFN2, per cell (iPSCs) or soma region (iNeurons) at different time points. Tether expression data from reference [42]. (**F**, **G**) Multispectral images and the corresponding line scan quantification of the pairwise contact between ER and peroxisomes, in the soma region of iNeurons at day7 and day14. Scale: 15 µm. (**H**) Plasmalogen biosynthesis pathway, illustrating how peroxisome-ER contacts are required for alkyl (O) precursors generated in peroxisomes to be transferred to the ER and converted into their final alkenyl (P) forms. (**I**) Fraction of the ether lipid species, normalized to the total level of each lipid species.

As iNeurons matured, we observed a subsequent increase in ER-lysosome, ER-peroxisome and ER-Golgi contacts (**Fig. 4D**). Similar dynamics were also observed when using absolute organelle volumes, without normalization to cell or soma volume (**Fig. S9B**), and when analyzing the fraction of organelle volume engaged in contacts (**Fig. S9C**). In neurites, the overall trend was preserved, but with fewer Golgi contacts and more lipid droplet contacts (**Fig. S9D, E**). These contacts all play roles in lipid metabolism: ER-peroxisome contacts are required for plasmalogen synthesis, while ER-Golgi and ER-lysosome contacts mediate cholesterol trafficking [35–37]. Both plasmalogens and cholesterol are enriched at synapses and in synaptic vesicles [38–40], consistent with the increase of these contacts at day 14. This is further supported by the additional increase in lipid droplet-organelle contacts in neurites, suggesting enhanced fatty acid trafficking for beta-oxidation and rapid membrane synthesis during neurite remodeling [41].

Because organelle contacts are mediated by protein tethers, we next examined how tether expression [42] relates to contact formation. The number of contacts for most organelle pairs correlated positively with the expression of their known tethers, with ER-peroxisome, mitochondria-peroxisome, and ER-mitochondria contacts ranking among the strongest associations (**Fig.4E-G**; **Fig. S10**). These correlations support a model in which the ER, mitochondria and peroxisomes play central, sequential roles during neuronal differentiation: an early phase of mitochondrial metabolic remodeling coordinated by ER-mitochondria and mitochondria-lysosome contacts; a concomitant increase in mitochondrial activity that necessitates augmented antioxidant defense, likely supported by peroxisome contacts [34]; and a later phase characterized by enhanced ER contacts with peroxisomes, Golgi and lysosomes, to remodel lipids required for membrane stability and synaptic homeostasis.

We further validated the expression levels of the protein tethers by Western blot (**Fig. S11A, B**) and investigated the number and volumes of ER-peroxisome contacts using AiryScan super-resolution microscopy, confirming an increase from day 7 to day14 (**Fig. S11C-E**). ER-peroxisome contacts mediated by VAPB at the ER and ACBD5 at peroxisomes are key hubs for plasmalogen biosynthesis, a class of membrane curvature-inducing lipids enriched at neuronal synapses [36, 38, 43, 44]. Plasmalogens are ether phospholipids whose synthesis begins in peroxisomes, where the alkyl (O) form is generated, and is completed in the ER, where the alkenyl (P) plasmalogen species is produced before being distributed to membranes (**Fig. 4H**). Consistent with increased engagement of this pathway, we observed an increase in the ether lipid fraction and in the alkenyl (P) and alkyl (O) group ratio during neuronal differentiation (**Fig. 4I**, **Fig. S12A, B; Data S8**), indicating an overall upregulation of plasmalogen synthesis and suggesting enhanced functional coupling between peroxisomes and the ER.

### ER-peroxisome contacts are a key feature of neuronal maturation

To test the functional role of ER-peroxisome contacts in neuronal differentiation, we next downregulated VAPB and ACBD5 and measured plasmalogen production (**Fig. 5A, B**; **Fig. S12C-G; Data S8**). Knockdown of VAPB and ACBD5, either alone or in combination, led to a marked reduction in total plasmalogen levels and in the plasmalogen fraction within major phospholipid classes (**Fig. 5C, D**; **Fig.S12F, G**) as previously reported [36]. The final alkenyl plasmalogen species (P) were most strongly reduced, indicating a defect in the terminal steps of plasmalogen biosynthesis that depend on intact ER-peroxisome tethering and lipid exchange (**Fig. 5C, E, Fig.S12F**). In addition, we observed a broad dysregulation of other major phospholipid classes, including PE, PC, PI and PA, upon VAPB and ACBD5 knockdown (**Fig. 5C**; **Fig. S12E**), consistent with a profoundly altered membrane lipid composition, as previously described [36].

**Figure 5:**
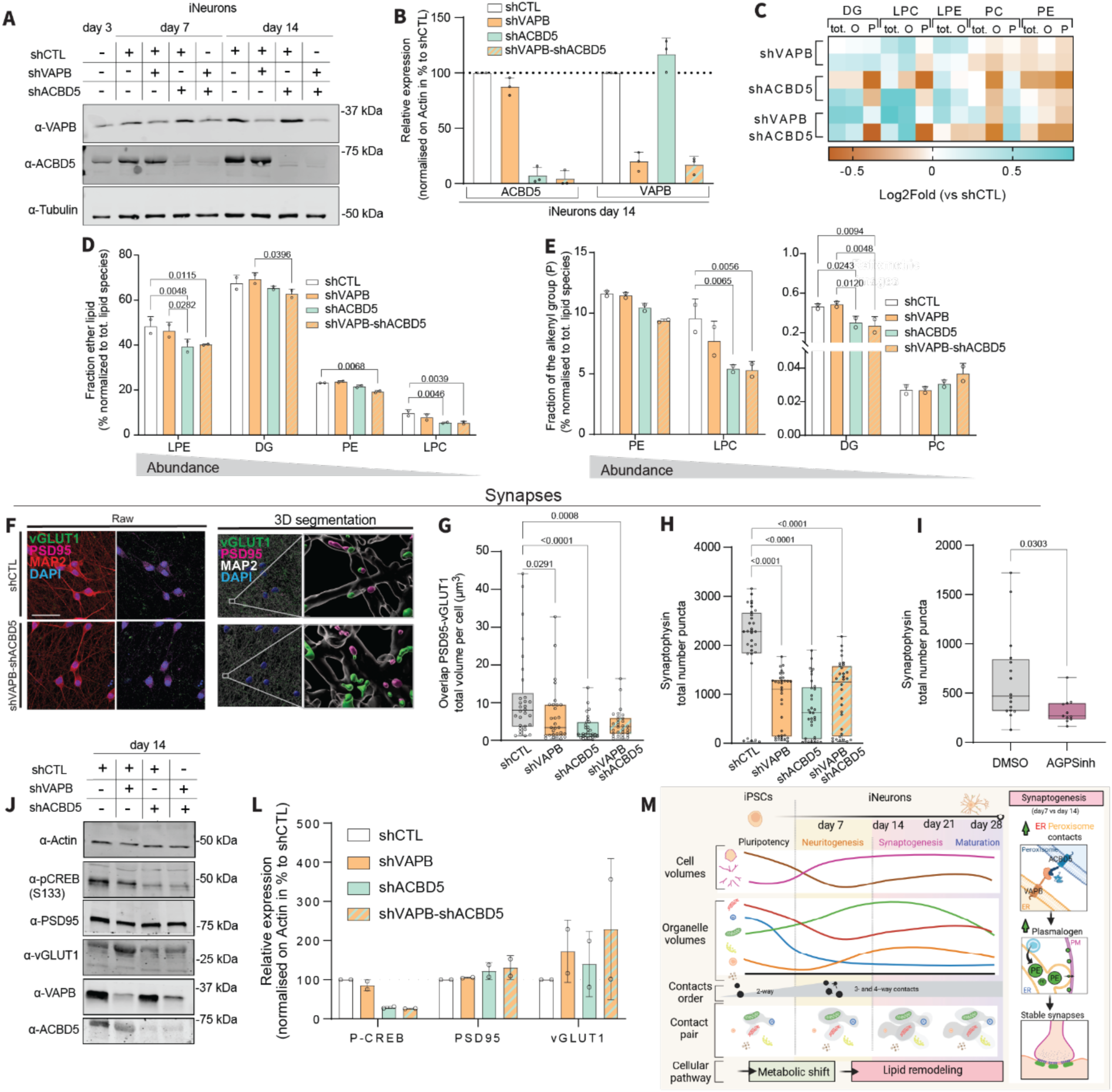
ER-peroxisome contacts are necessary for stable synapses. (**A**) Immunoblot and its (**B**) quantification of the tethering proteins VAPB and ACBD5 in iNeurons at day3, 7, and 14, after knock-down of VAPB and ACBD5, alone or in combination. (**C**) Total levels (tot), alkyl (O), and alkelyl (P) forms of different classes of lipids. (**D**) Fraction of the ether lipid species, normalized to the total level of each lipid species. (**E**) Fraction of the alkenyl (P) group, normalized to the total level of each lipid species. (**F**) Immunofluorescence of pre-synaptic (vGLUT1) and post-synaptic (PSD95) markers, together with a neuronal marker (MAP2) and nuclei (DAPI) in iNeurons at day 14, after knockdown of VAPB and ACBD5 or in the corresponding control (Ctl). Scale: 15 µm. (**G**) Quantification of the overlap between vGLUT1 and PSD95 puncta in iNeurons at day14, after knock-down of VAPB and ACBD5, individually or in combination. (**H**) Quantification of the number of synaptophysin positive puncta in iNeurons at day 14, after knock-down of VAPB and ACBD5, individually or in combination, (**I**) or after inhibition of the AGPS enzyme. (**J**, **L**) Immunoblot and its quantification, of synaptic markers (PSD95 and vGLUT1), tethering proteins (VAPB and ACBD5), together with phosphoCREB (S133) in iNeurons at day3, 7, and 14, after knock-down of VAPB and ACBD5, individually or in combination. (**M**) Graphical abstract showing major changes in organelle morphology and contacts throughout neuronal differentiation. Two-way ANOVA; p-value versus control knockdown.

The lipid composition of membranes and especially at the synapses is extremely tightly regulated in order to provide both plasticity and structural stability [43], and plasmalogen species, such as PE and LPC, play a crucial role in conferring membrane curvature, vesicle fusion and platforms for synaptic protein complexes. These lipids are particularly enriched at pre-and post-synaptic membranes [38, 39, 44], where they support docking and function of key synaptic proteins. We therefore asked whether synapses are compromised when ER-peroxisome contact were reduced. Immunostaining of PSD-95 and vGLUT1 (**Fig. 5F, G**; **Fig. S13A-E**), as well as synaptophysin (**Fig. 5H**; **Fig. 13F, G**), revealed decreased synapses density, with no major change in soma or neurites volumes when VAPB and ACBD5 were knocked down (**Fig. S13A, F**). The reduction in individual vGLUT1 puncta was particularly pronounced, consistent with defects in synaptic vesicle trafficking and release, processes that depend critically on the lipid composition and biophysical properties of presynaptic and vesicular membranes (**Fig. S13B, C-D**). To independently perturb plasmalogen synthesis, we treated iNeurons with an inhibitor of alkylglyceronephosphate synthase (AGPS), a peroxisomal enzyme that catalyzes one of the first steps of ether lipid and plasmalogen biosynthesis. AGPS inhibition phenocopied the synaptic defects observed upon VAPB and ACBD5 knockdown (**Fig. 5I**; **Fig. S13H-J**), further linking impaired plasmalogen production to synaptic dysfunction. These dysregulated synaptic structures resulted in reduced neuronal activity as indicated by decreased levels of phosphorylated-CREB, the upstream regulator of c-Fos, when VAPB and/or ACBD5 were depleted (**Fig. 5J, L**). Together, these findings identify ER-peroxisome contacts as a key regulator of plasmalogen-dependent membrane homeostasis and synaptic maturation in neurons.

## Discussion

Previous research has established that organelles undergo significant remodeling during neuronal differentiation [1, 3–6, 8]. This study provides a temporal map of organelle network rewiring, revealing that neuronal differentiation is accompanied by a tightly timed and compartment-specific remodeling of the organelle system. Using MSI of eight organelles, quantitative organelle signature analysis, and lipidomics in hNGN2-driven human neurons, we find that organelles do not simply increase or decrease in bulk: they rescale with cell geometry, change size, number and distribution in soma and neurites, and adopt compartment-specific identities that together establish a neuron-specific organelle landscape (**Fig. 5M**).

At the level of interactions, we reveal a progressive increase in organelle connectivity and higher-order contacts as neurons differentiate. We observed first an increase in the contact number, followed by contact volume, indicating that the organelle network is first laid out and then strengthened. Mitochondria emerge as an early hub, with increased ER-mitochondria, mitochondria-lysosome and mitochondria-peroxisome contacts, consistent with extensive remodeling of the mitochondrial interactome to support metabolic rewiring at the onset of neuronal differentiation. These observations align with existing knowledge about shifts in mitochondrial composition, form, and function throughout neuronal differentiation [7–9]. At later stages, contacts of the ER with peroxisomes, lysosomes and Golgi suggest a shift toward lipid remodeling and membrane specialization as neurons mature (**Fig. 5M**).

Within this framework, ER-peroxisome contacts act as a key node linking organelle rewiring to synapse formation. ER-peroxisome contacts have previously been implicated in plasmalogen biosynthesis in non-neuronal cells [36]. While synapses and synaptic vesicles are known to be enriched in plasmalogens [38, 39], the role of ER-peroxisome contacts in neuronal membrane homeostasis and synapse formation was unappreciated. We show that these contacts support plasmalogen biosynthesis and broader phospholipid homeostasis throughout neuronal differentiation; their disruption via VAPB/ACBD5 knockdown or AGPS inhibition reduces plasmalogens, and impairs synaptic organization, with decreased PSD-95, vGLUT1 and synaptophysin puncta and reduced CREB activation. Together, these findings identify ER-peroxisome contacts as a critical feature of neuronal maturation. Moreover, mutations in the ER-peroxisome tether VAPB cause a rare familial form of amyotrophic lateral sclerosis (ALS8), which is associated with early synaptic dysfunction [45, 46]. While several studies have emphasized synaptic mitochondrial impairment in ALS, our data support the idea that disrupted ER-peroxisome coupling and the resulting defects in plasmalogen-dependent membrane homeostasis may represent a parallel, and potentially targetable, mechanism contributing to synaptic vulnerability [47]. Consistent with this model, reduced plasmalogen abundance has been reported as a hallmark of ALS and other neurodegenerative disorders, including Alzheimer’s disease, further highlighting ER-peroxisome contacts as a candidate druggable axis [48, 49]. Overall, our study illustrates how MSI of the organelle interactome can be used to follow cell state trajectories and uncover disease-relevant vulnerabilities.

## Materials and Methods

### Cells and culture conditions

KOLF2.1J wild-type (KOLF2.1 wt) human induced pluripotent stem cells (iPSCs) (17) were obtained from Dr. Bill Skarnes (The Jackson Laboratory) and were used to generate an iPSC piggyBac tetracycline-on human neurogenin-2 (iPSC-PB-TO-hNGN2) cell line. Cells were incubated at 37 DC, 5% CO2 and cultured under feeder-free conditions in StemFlex medium (Gibco A3349401) on Vitronectin (Vitronectin (VTN-N) Recombinant Human Protein, Truncated, Gibco A14700). The cells were passaged using ReLeSR (STEMCELL Technologies) in the absence of ROCK inhibitor unless thawing or making frozen stocks.

### Generation of PB-TO-hNGN2-iPSCs

An iPSC-PB-TO-hNGN2 stable cell line was generated as previously described [17]. Briefly, KOLF2.1 wt iPSCs were transfected with piggyBac plasmids encoding rtTA and an NGN2-Puro cassette (plasmids kindly provided by Dr. Michael Ward, National Institute of Neurological Disorders and Stroke). Two transfection conditions were used: 0.5 µg total piggyBac DNA (clone 1) or 1.0 µg (clone 2), for a well of a 6-well plate. Following transfection, cells were selected for stable integration with puromycin (0.5 µg/mL) and expanded as non-clonal pools. Clones were evaluated for proliferation and differentiation capacity using the sulforhodamine B (SRB) assay, and pluripotency marker expression was validated by immunofluorescence, western blotting, and RT-qPCR. Clone 1 showed the most robust proliferation, pluripotency marker expression, and differentiation efficiency and was therefore used for downstream experiments.

### Differentiation into cortical neurons

iPSC-PB-TO-hNGN2 cells were differentiated into cortical neurons as previously described [17] with a modification. In brief, iPSCs were seeded dissociated on vitronectin coated plates, in induction medium [DMEM/F12 with HEPES (Gibco, 11330032); N2 supplement 100X (Gibco, 17502048); non-essential amino acids (NEAA) 100X (Gibco, 11140050) and supplemented with ROCK inhibitor 1X (RevitaCell supplement 100X, Gibco A2644501) and doxycycline at a final concentration of 0.5 µg/mL (Sigma, D9891)]. The next day medium was replaced with induction medium without ROCK inhibitor and with fresh doxycycline (1 µg/mL). After 2 days, the pre-induced iPSCs were passaged in accutase (StemPro™ Accutase™ Cell Dissociation Reagent, Gibco A1110501) and seeded single cell (43,000 per well) on poly-L-ornithine-(Sigma, P3655; 10x stock: 50 mg in 50 ml borate buffer) and laminin- 10 µg/ml (Gibco, 23017015) coated plates (Thermo Scientific™ Nunc™ Lab-Tek™ II Chambered Coverglass, 8-well # 155409) using induction medium supplemented with doxycycline at a final concentration of 0.5 µg/mL. The next day of the seeding the medium was changed to cortical neuron culture medium (CM) [BrainPhys neuronal medium without phenol red (STEMCELL Technologies, 05791); SM1 supplement, 50X (STEMCELL Technologies, 05711); BDNF (10 µg/ml) in PBS containing 0.1% IgG and protease-free BSA (PeproTech 450-02); NT-3 (10 µg/ml) in PBS containing 0.1% IgG and protease-free BSA (PeproTech 450-03); laminin final concentration 1 µg/ml (Gibco 23017015)], and freshly supplemented with doxycycline at a final concentration of 0.5 µg/mL. The iPSC-derived neurons (iNeurons) were kept for 28-30 days with half media changes at least once per week with freshly prepared CM. We further tested the successful differentiation into cortical neurons by staining for different neuronal markers.

### iNeuron transduction

Lentiviral short hairpin RNA (shRNA) expression plasmids were purchased from Sigma (MISSION® TRC1 pLKO.1-puro): a non-mammalian shRNA control (shCTL), TRCN0000152239 targeting VAPB (shVAPB), and TRCN0000107026 targeting ACBD5 (shACBD5). Lentivirus was concentrated using Lenti-X Concentrator (Takara Bio 631231) and resuspended in DPBS. Viral titers were estimated by RT-qPCR (abm; qPCR Lentivirus Titer Kit, LV900) and viral stocks were diluted with DPBS to equalize the theoretical viral concentration across conditions. On the day of transduction, half of the culture medium was collected and iNeurons (day 4) were incubated with diluted lentivirus (1:20) in fresh cortical medium. After 24 hours, medium was completely replaced with a 1:1 mix of fresh cortical neuron medium and the previously collected conditioned medium. To control for total viral/shRNA load and minimize potential competition effects, single knockdowns were performed by mixing each targeting virus 1:1 with control virus: shVAPB = shVAPB + shCTL; shACBD5 = shACBD5 + shCTL. Controls received shCTL + shCTL, and double knockdown received shVAPB + shACBD5.

### AGPS inhibitor treatment

Pre-induced iNeurons (120,000 cells per well) were seeded onto 12-mm diameter #1.5H thick precision (Electron Microscopy Sciences German Glass Coverslips #1.5, 12 mm; 72290-04), and cultured as described above. The alkylglycerone phosphate synthase (AGPS) inhibitor, AGPS-IN-2i (AOBIOUS; AOB17269), was prepared and used following the manufactures instruction. Briefly, AGPS inhibitor stock was made in DMSO at 50 mM, and used within one month, at a final concentration of 250 µM in cortical medium. At day 10 of differentiation, 250 µM AGPS inhibitor or corresponding 0.005% of DMSO as control were added to the wells. Every second day, a half media change was performed and AGPS inhibitor/DMSO were replaced freshly in the media, until day 14, when samples were fixed and processed as described below.

### Transfection of iPSCs

The cells were dissociated and seeded (6,000 per well) on vitronectin coated plates (Thermo Scientific™ Nunc™ Lab-Tek™ II Chambered Coverglass, 8-well # 155409) in StemFlex media supplemented with ROCK inhibitor at a final concentration of DBP (RevitaCell supplement 100X, Gibco A2644501). The day after seeding, the medium was replaced with StemFlex without ROCK inhibitor, and the cells were left to grow for 4 days. The medium was replaced with Essential 8 (Gibco, A1517001; not supplemented) approximately 20 minutes before transfection. The transfection mix was prepared in Opti-MEM (Gibco 31985070) with 0.5 μl of Lipofectamine Stem Reagent (Invitrogen, STEM00001), and 500 ng of total DNA divided as follows: 167 ng of pEIF1a::Transposase (gifted by Dr. Michael Ward), 150 ng of pEIF1a::LAMP1::mTurquoise (lysosomes), 100 ng of pEIF1a::Cox8(1-26)::eGFP (mitochondria), 40 ng of pEIF1a::Sit::OxVenus (Golgi), 15 ng of pEIF1a::mOrange2::SKL (peroxisomes), and 150 ng of pEIF1a::Sec61β::mApple (endoplasmic reticulum) (Twist Technologies). The mix was applied dropwise and left incubating for four hours at 37 DC. The medium was then changed back to StemFlex.

### Transfection of iNeurons

At each time point along the differentiation (day6, 13, 20, and 27), the iNeurons were transfected as previously described [10, 14, 15, 50], with some modifications. The transfection mix was prepared in Neurobasal (Gibco 21103049) with 1µl of Lipofectamine2000 (Invitrogen, 11668019), and 1 µg of total DNA divided as follows: 167 ng of pEIF1a::Transposase (gifted by Dr. Michael Ward), 120 ng of pEIF1a::LAMP1::mTurquoise, 120 ng of pEIF1a::Cox8(1-26)::eGFP, 60 ng of pEIF1a::SiT::OxVenus, 20 ng of pEIF1a::SKL::mOrange, and 120 ng of pEIF1a::Sec61β::mApple (Twist Technologies). The medium of iNeurons was changed to Neurobasal (without glutamine) and pre-incubated for at least 30 minutes before adding the transfection mix. Next, half of the medium was removed, and the transfection mix was added dropwise. After 1 hour of incubation, the medium was completely replaced with BrainPhys Imaging Optimized Medium (STEMCELL Technologies, 05796) supplemented with SM1 supplement, 50X (STEMCELL Technologies, 05711); BDNF (10 µg/ml) in PBS containing 0.1% IgG and protease-free BSA (PeproTech 450-02); NT-3 (10 µg/ml) in PBS containing 0.1% IgG and protease-free BSA (PeproTech 450-03); Laminin final concentration 1 µg/ml (Gibco 23017015)].

### Labeling and microscopy of live iPSCs and iNeurons

24 h after transfection, the iNeurons were tested for vitality by NeuroFluo at a final concentration of 0.20 µM (STEMCELL Technologies, 01801); staining data not shown. iPSCs and iNeurons were labelled with Lipi-Blue dye (Dojindo) and CellMask Far-Red dye (Invitrogen, C10046) for at least 20 minutes and 10 minutes, respectively, prior to imaging. Images were acquired on a Zeiss 880 laser scanning confocal microscope equipped with a 32-channel multi-anode spectral detector (Carl Zeiss) in lambda mode at 8.9 nm bins (collecting wavelengths 410-695), with 63x/1.4 NA objective lens, and a 2.2x zoom. All fluorophores were excited simultaneously using 405, 458, 514, 594, and 633 nm lasers, with a 458/514/561/633 nm main beam splitter. Z-stacks were acquired with 1.2 μm slices, at a scan speed of 1.97 s/frame. Live imaging was performed with stable 5% CO2 at 37 □C, and each imaging session was not longer than 1-2 hours. Z-stacks spanning the entire cell were acquired in lambda mode (lambda scan 410-695 nm, 8.9 nm resolution as described above) and subsequently processed by linear unmixing to produce 10 distinct channels using ZEN Blue 2.3 software (Carl Zeiss). Channels 0-8, correspond to each of the fluorescently labeled structures: lipid droplets, nuclei, lysosomes, mitochondria, Golgi, peroxisomes, endoplasmic reticulum (ER), and plasma membrane; channel 9 is the residual signal from the unmixing process, and channel 10 is the transmitted brightfield. Linear-unmixed images were subsequently deconvolved using Huygens Essential software version 23.04 (Scientific Volume Imaging, The Netherlands, http://svi.nl). For each time point, one image was chosen to generate a template for batch processing. Excitation and emission maxima (Ex/Em max) for each fluorophore were estimated from the metadata, with assisted pinhole adjustment; deconvolution was performed with the standard classic maximum likelihood estimation (MLE) algorithm, no acuity adjustment. Different deconvolution settings were applied for each channel, with signal-to-noise ratios (SNRs) ranging from 6 to 11. Other settings were implemented as follows: lipid droplets (lowest; radius 0.7µm), nuclei (lowest; radius 2 µm); lysosomes (inner-near; radius 0.7 µm); mitochondria (inner-near; radius 0.7 µm); Golgi (inner-near; radius 0.7 µm); peroxisomes (inner-near; radius 0.5 µm); ER (lowest; radius 0.3 µm); plasma membrane (lowest; radius 0.3 µm). The multispectral microscopy dataset is available at BioImage Archive accession number (S-BIAD2983) (https://www.ebi.ac.uk/bioimage-archive/studies), and is composed of z-stacks of 43 undifferentiated iPSCs (from 5 biological replicates); 97 iNeurons day7 and 94 iNeurons day14 (from 4 biological replicates); 82 iNeurons day21 and 79 iNeurons day28 (from 5 biological replicates).

Brightfield images were acquired using an EVOS M5000 inverted microscope (Thermo Fisher Scientific) equipped with a transmitted-light illumination module using a 20x objective.

### Coverslip preparation and fixation of iNeurons

German glass coverslips (12 mm diameter, #1.5H; Electron Microscopy Sciences, 72290-04) were sonicated at 42 kHz in 70% ethanol for 30 min, repeated three times with fresh 70% ethanol. Coverslips were air-dried in a biosafety cabinet, placed in 24-well tissue culture plates (Fisherbrand™, FB012929), and UV-sterilized for 15 min. For iNeuron cultures, coverslips were coated with 0.1 mg/mL poly-L-ornithine hydrobromide (Sigma, P3655) in borate buffer (10 mM boric acid (Sigma B6768), 2.5 mM sodium tetraborate (Sigma 221732), 7.5 mM sodium chloride (Sigma S7653), and 0.1 M sodium hydroxide (Sigma 71463)) for 2 days at 37 °C. Coverslips were washed three times with autoclaved Milli-Q water and then coated with laminin (10 µg/mL; Gibco, 23-017-015) in cold DPBS for 2 hours at 37 °C.

iNeurons were cultured as described above and fixed at the indicated time points by adding pre-warmed 8% paraformaldehyde (PFA) directly to the culture medium to reach a final concentration of 4%, followed by incubation for 15 min at 37 °C. Fixed neurons were gently washed three times with DPBS (fully submerged to prevent drying) and stored at 4 °C, protected from light. Samples were used within one month.

### Immunostaining and imaging of iPSC pluripotency markers

Dissociated iPSCs (6,000 cells per well) were seeded onto vitronectin-coated 8-well chambered coverglass (Thermo Scientific™ Nunc™ Lab-Tek™ II, #155409) in StemFlex medium supplemented with ROCK inhibitor (RevitaCell, 1X final; Gibco A2644501). The following day, medium was replaced with StemFlex without ROCK inhibitor, and cells were cultured for 4 days. Cells were washed three times with 1x DPBS and fixed with pre-warmed 4% PFA in DPBS for 15 min at 37 °C. After three washes in DPBS, cells were blocked and permeabilized for 1 h at room temperature in 4% BSA and 0.3% Triton X-100 in DPBS. Cells were incubated overnight at 4 °C with primary antibodies (300 µL per well): rabbit anti-OCT4 (1:300, Abcam ab181557) and mouse anti-SSEA4 (1:300, ThermoFisher mc813-70). The next day, cells were washed three times with DPBS and incubated for 1 h at room temperature in the dark with species-appropriate secondary antibodies (1:1000) diluted in blocking solution: anti-rabbit Alexa Fluor 488 (abcam; ab150073); anti-mouse Alexa Fluor 568 (Invitrogen; A10037).

For wild-type iPSCs, nuclei were stained with DAPI (1:10,000; Thermo Scientific 62248) for 10 min in DPBS prior to imaging. Cells were imaged in 200 µL DPBS per well. Images were acquired on a Zeiss LSM 800 laser scanning confocal microscope using a 63x/1.4 NA oil-immersion objective at 1.2x zoom.

### Immunostaining and imaging of iNeuron differentiation markers

Cells were cultured and fixed at the indicated time points (days 3, 7, 14, 21 and 28) as described above. Once all time points were collected, coverslips were processed in parallel. Coverslips were blocked and permeabilized for 1 h at room temperature in 4% BSA and 0.3% Triton X-100 in DPBS. Coverslips were next incubated overnight at 4 °C with mouse anti- ankyrin-G N106/36 (1:100; antibodies Inc SKU: 75-146); chicken anti-MAP2 (1:1000, Invitrogen PA1-10005); rabbit anti-β-III-Tubulin (abcam; ab18207). The next day, coverslips were washed three times and incubated for 1 h at room temperature in the dark with secondary antibodies diluted in blocking solution: anti-Rabbit AF568 (1:1000, Invitrogen; A10042); anti-Mouse AF488 (1:1000, Invitrogen A21202); anti-Chicken AF647 (1:1000, Invitrogen A-21449). Coverslips were washed and mounted using ProLong™ Diamond Antifade mountant with DAPI (Invitrogen, P36971) and allowed to cure for 48 h before imaging.

### Immunostaining, Fast Airyscan imaging and analysis

Cells were cultured and fixed at the indicated time points (iPSCs; iNeurons days 3, 7, 14 and 21) as described above. Once all time points were collected, coverslips were processed in parallel. Coverslips were blocked and permeabilized for 1 h at room temperature in 4% BSA and 0.3% Triton X-100 in DPBS. Coverslips were first incubated overnight at 4 °C with rat anti-TOM20 (1:100, Abcam ab289670) in a humid chamber. The following day, coverslips were washed three times with DPBS and incubated for 1 h at room temperature in the dark with anti-rat Alexa Fluor 568 (1:1000, Thermo Fisher A78946) diluted in blocking solution. Coverslips were then washed three times with DPBS and incubated overnight at 4 °C with mouse anti-PMP70 (1:300, Sigma SAB4200181), rabbit anti-LAMP1 (1:300, Cell Signaling #9091), and goat anti-calnexin (C-terminal) (1:100, Abcam ab219644). The next day, coverslips were washed three times and incubated for 1 h at room temperature in the dark with secondary antibodies diluted in blocking solution: anti-mouse Alexa Fluor 488 (1:1000, Thermo Fisher A32787), anti-goat Alexa Fluor 647 (10 µg/mL, Thermo Fisher A21447), and anti-rabbit Alexa Fluor 405 (1:500, Abcam ab175651). Coverslips were washed and mounted using ProLong™ Glass Antifade mountant and allowed to cure for 48 h before imaging. Z-stacks spanning the entire cell were acquired on a Zeiss LSM 880 laser scanning microscope using a 63x/1.4 NA oil-immersion objective at 2.4x zoom in Fast Airyscan mode. Images were processed using 3D Airyscan in ZEN Black 2.3 and subsequently deconvolved in ZEN Blue 2.3 (Carl Zeiss).

Subsequent segmentation of organelles and soma-neurite compartments were performed using InferSubC v2 beta (v2.0.0b1; https://github.com/SCohenLab/infer-subc/tree/v2.0.0b1). Quantification was performed per cell (soma region) for iNeurons and per field-of-view for iPSCs. For iPSCs, whole-field nuclei were manually counted and used to normalize organelle counts and total volumes to a per-cell basis.

### Immunostaining, imaging and analysis of synapses

Pre-induced iNeurons were seeded onto 12-mm diameter #1.5H precision glass coverslips (Electron Microscopy Sciences, 72290-04), cultured and transduced as described above. At day 14, cells were fixed, washed three times with DPBS, and blocked and permeabilized for 1 h at room temperature in 4% BSA and 0.3% Triton X-100 in DPBS.

For synaptic marker staining, coverslips were incubated overnight at 4 °C with primary antibodies diluted in blocking solution. For excitatory synapses, primary antibodies were rabbit anti-PSD95 (1:300, Abcam ab18258), mouse anti-vGLUT1 (1:300, Sigma MAB5502), and chicken anti-MAP2 (1:1000, Invitrogen PA1-10005). For presynaptic marker analysis, primary antibodies were rabbit anti-Synaptophysin-1 (1:300, Synaptic Systems 101 002) and mouse anti-βIII-Tubulin (1:300, Abcam ab78078). After three washes in DPBS, coverslips were incubated for 1 h at room temperature in the dark with species-appropriate Alexa Fluor-conjugated secondary antibodies (see below), diluted in blocking solution. Coverslips were washed and mounted using ProLong™ Diamond Antifade mountant with DAPI (Invitrogen, P36971) and allowed to cure for 48 h before imaging.

For excitatory synapses, secondary antibodies were: anti-Rabbit AF647 (Invitrogen; A32795TR); anti-Mouse AF488 (Invitrogen; A21202); anti-Chicken AF594 (Invitrogen; A78951). For presynaptic marker analysis, secondary antibodies were: anti-Rabbit AF488 (abcam; ab150073); anti-Mouse AF568 (Invitrogen; A10037). Z-stack images spanning the entire cell were acquired on a Zeiss LSM 800 laser scanning confocal microscope using a 63x/1.4 NA oil-immersion objective at 1.4x digital zoom. Identical acquisition settings were used across conditions within each experiment.

Volumetric synapse quantification was performed using Imaris software (version 10.2.0; Bitplane). Briefly, nuclei, neuronal processes, and synaptic puncta were segmented using intensity-based surface detection with local background subtraction. Neuronal morphology was defined using MAP2 or β-III-Tubulin surfaces, and synaptic objects (PSD95, vGLUT1 or synaptophysin) were segmented with size and distance constraints relative to neuronal surfaces to restrict detection to synaptic regions. Synapses were defined based on the presence and spatial overlap of pre- and postsynaptic markers or by presynaptic puncta density, depending on the experiment. Volumes, counts and overlap measurements were extracted from Imaris and used for quantitative analyses and graph generation.

### RNA extraction and RT-qPCR

iPSCs (80,000 cells per well) and pre-induced iNeurons (650,000 cells per well) were seeded in 6-well plates (Fisherbrand™ Surface Treated Sterile Tissue Culture Plates; FB012927) and cultured as described above. iPSCs were collected after 4 days of growth, while iNeurons were collected at the indicated differentiation time points (days 7, 14, 21, and 30). At collection, medium was aspirated, cells were washed once with DPBS, and lysed directly in TRIzol reagent (500 µL per well; Invitrogen, TRIzol™ Reagent 15596026). Lysates were stored at −80 °C until processing. Once all time points were collected, RNA was extracted according to the manufacturer’s instructions using acid phenol:chloroform (500 µL; pH 4.5, with IAA, 125:24:1; Invitrogen, AM9720), with the addition of 5 µg glycogen (Thermo Scientific™, FERR0561) as a carrier. RNA pellets were resuspended in 30 µL DEPC-treated water. RNA purity and integrity were assessed using 1.2% agarose gel electrophoresis and spectrophotometry (NanoDrop; A260/280 and A260/230 ratios).

cDNA was performed using 1 µg of total RNA for each sample using Applied Biosystems High-Capacity cDNA reverse transcription kit as per manufacturer’s instructions. Quantitative real-time PCR (RT-qPCR) was performed using SYBR Green according to the manufacturer’s guidelines. Each sample was run in technical duplicate, with two independent biological replicates per time point. β-Actin was used as an endogenous control and samples were assessed for the expression of pluripotent genes (OCT4, SOX2, NANOG) and neuronal genes (vGLUT1, PSD95, and SYN1). Relative gene expression levels were calculated using the comparative Ct (Livak) method. Briefly, raw Ct values were averaged across technical duplicates for each target gene and normalized to β-Actin within each biological replicate to obtain ΔCt values. The mean ΔCt of iPSC biological replicates was used as the reference condition to calculate ΔΔCt values for each differentiation time point. Data are presented as fold changes relative to iPSCs, calculated as 2^−ΔΔCt.

### Lipidomics sample preparation, acquisition, and analysis

#### Cell collection and sample preparation

iPSCs and iNeurons were cultured in 100-mm dishes as described above. Cells were collected at the indicated differentiation time points (iPSCs; iNeurons at day 3, day 7, and day 14), as well as at day 14 following lentiviral knockdown of VAPB and/or ACBD5. At collection, cells were washed once with DPBS and incubated with Accutase (Sigma; A6964) for 10 min at 37 °C. Cells were washed once with cold DPBS pelleted by centrifugation for 10 min at 13,000 rpm at 4°C, and the supernatant was carefully aspirated. Cell pellets were flash-frozen in liquid nitrogen and stored at −80 °C until extraction. Lipid extraction. For lipid extraction, 1 mL of MTBE was added to each cell pellet, samples were vortexed and transferred to Eppendorf tubes. Methanol (300 µL) containing internal standards was added, and samples were shaken for 10 min. Phase separation was induced by adding 200 µL of water, followed by centrifugation at 2,000 rcf for 10 min. The upper organic phase was collected, dried under vacuum, and reconstituted in 100 µL of isopropanol (IPA) for LC-MS analysis. Avanti EquiSplash deuterated lipid mix was used as an internal standard and spiked into the methanol at 1.5 µg/mL prior to extraction.

#### LC-MS acquisition

Lipidomic analysis was performed using a Thermo Q Exactive Plus mass spectrometer coupled to a Waters Acquity H-Class UPLC system. Lipids were separated on a 100 x 2.1 mm, 2.1 µm Waters BEH C18 column. Mobile phases consisted of A: 60/40 acetonitrile/water and B: 90/10 isopropanol/acetonitrile, both supplemented with 10 mM ammonium formate and 0.1% formic acid. The flow rate was 0.2 mL/min. The gradient started at 32% B, increased to 40% B at 1 min (held until 1.5 min), 45% B at 4 min, 50% B at 5 min, 60% B at 8 min, 70% B at 11 min, and 80% B at 14 min (held until 16 min), before returning to 32% B and equilibrating for 4 min. Data were acquired in positive/negative ion-switching mode using top-5 data-dependent MS/MS fragmentation.

#### Lipid identification and quantification

Raw data were processed using LipidSearch software. Lipids were identified based on MS/MS fragmentation with a precursor mass tolerance of 5 ppm and a product ion mass tolerance of 8 ppm. Identifications were generated for individual samples and subsequently aligned across experimental groups. Quantification was performed using EquiSplash internal standards, and processed data were exported to Excel for downstream analysis.

#### Plasmalogen analysis and normalization

Protein content was measured for each sample. Lipid abundances were first normalized to internal class standards (ICS) for each lipid species and subsequently normalized to protein content where indicated. For differentiation analyses, lipid abundances normalized to ICS alone were reported, to avoid inflation effects due to increased neurite mass during neuronal maturation. For VAPB and ACBD5 knockdown experiments in iNeurons (day 14), lipid abundances were normalized to both ICS and protein content.

## Western blotting

Dissociated iPSCs (80,000 cells per well) and iNeurons (650,000 cells per well) were cultured as described above in 6-well plates (Fisherbrand™, FB012927) coated with vitronectin or poly-L-ornithine/laminin, respectively. Cells were lysed on ice in RIPA buffer (50 mM Tris-HCl pH 8.0, 150 mM NaCl, 1% NP-40, 0.5% deoxycholate, 0.1% SDS) supplemented with 15 mM sodium pyrophosphate, 50 mM sodium fluoride, 40 mM β-glycerophosphate, 1 mM sodium orthovanadate, PMSF, leupeptin and aprotinin, as well as protease and phosphatase inhibitor cocktails (Protease inhibitor cocktail, Sigma P8340; Phosphatase inhibitor cocktails 2 and 3, Sigma P5726 and P0044; each at 1:100). Adherent cells were scraped, collected, and centrifuged at 13,000g for 5 min at 4 °C to obtain post-nuclear supernatants. Protein concentration was determined using the Pierce BCA assay (Thermo Scientific, 23225). Samples were mixed with 6X Laemmli buffer containing β-mercaptoethanol and denatured for 10 min at 75 °C. For each sample, 50 µg protein was resolved by 12% SDS-PAGE and transferred to 0.2-µm nitrocellulose membranes (Bio-Rad 162-0112) by wet transfer overnight at 4 °C at constant current (320 mA). Membranes were briefly rinsed in water and stained with Ponceau S (Thermo scientific A40000279) to verify transfer quality and uniform loading, then de-stained in TBST and blocked in 5% milk in TBST for 1 h at room temperature. Membranes were incubated overnight at 4 °C with primary antibodies diluted in 5% milk in TBST, washed three times for 10 min in TBST, and incubated with IRDye-conjugated secondary antibodies diluted in 5% milk in TBST for 1 h at room temperature in the dark. Membranes were washed three times for 5 min in TBST and imaged on an Odyssey CLx (LI-COR). Band intensities were quantified using Plot Profile in FIJI/ImageJ (https://imagej.net/software/fiji).

Primary antibodies used were: rabbit anti-OCT4 (1:1,000, abcam ab181557), rabbit anti-NANOG (1:500, abcam ab21624), rabbit anti-ACBD5 (1:1,000, Proteintech 21080-1-AP), rabbit anti-VAPB (1:1,000, Proteintech 14477-1-AP), mouse anti-α-tubulin DM1A (1:10,000, Abcam ab7291), rabbit anti-βIII-tubulin (1:1,000, Abcam ab18207), and mouse anti-actin SPM161 (1:5,000, Santa Cruz sc-56459). Secondary antibodies were IRDye anti-mouse 800CW (1:10,000, LI-COR 925-68074) and IRDye anti-rabbit 680RD (1:10,000, LI-COR 926-68073).

### Line scan quantification

Images were randomly selected and a single z-plane was used for analysis. A line region of interest (ROI) was drawn in FIJI to intersect the organelles of interest. ROIs were saved using the ROI Manager and used to extract fluorescence intensity profiles across all channels. The same ROI was applied across time points for the corresponding regions; when necessary, the ROI was rotated without changing its length. Intensity profiles were exported using the Plot Profile tool, and grey values were normalized and plotted as a percentage of the maximum intensity for each channel.

### Multispectral image analysis

#### 3D multispectral image segmentation

Instance segmentations of organelles were generated from independent intensity channels in deconvolved 3D z-stack images. Python-based segmentation workflows (1.2-1.7) from infer-subc (v2.0.0b1; https://github.com/SCohenLab/infer-subc/tree/v2.0.0b1) were used; methods are described in detail in the associated Jupyter Notebooks (https://github.com/SCohenLab/infer-subc/tree/v2.0.0b1/notebooks/part_1_segmentation_workflows). Lysosomes and mitochondria were the only organelles that underwent declumping to separate touching organelles using (“declumping”, https://github.com/SCohenLab/Extended-analysis-for-infer-subc-v2.0.0b1; now available as the final step in each plugin workflows). For iPSCs, whole-cell masks (CellMask) and nuclear masks were generated using infer-subc workflow 1.1d. For iNeurons, whole-cell masks were generated using infer-subc workflow 1.1c; however, due to silencing of the BFP2 nuclear reporter, an alternative approach was used to segment the nuclei that is based on the inverse of the ER signal (“Find Nuclei” and “Find Nuclei_Batch process”; https://github.com/SCohenLab/Extended-analysis-for-infer-subc-v2.0.0b1). When required, CellMasks were manually refined in Napari using plasma membrane and ER signals. For iNeurons, CellMasks were subsequently processed using an automated pipeline to separate soma and neurite regions (“soma_neurite_filter”, https://github.com/SCohenLab/Extended-analysis-for-infer-subc-v2.0.0b1; now available as workflow 1.8).

#### 3D multispectral organelle quantification

Quantitative analysis of 3D organelle organization was performed as previously described [10], with additional features including organelle declumping, segmentation of soma vs. neurites, and analysis of multi-way organelle interactions. Briefly, for each organelle and organelle interaction, morphological (number, volume, shape) and spatial distribution metrics were extracted. Organelle interactions were defined as regions of voxel-wise overlap between segmented organelle objects of different types.

Although the organelle segmentations were generated across the full image frame, all downstream analyses were restricted to soma and neurite regions for each cell (“subcompartments”) and batch-processed using the infer-subc organelle signature analysis pipeline. The analyses reported here were run using the infer-subc codebase at version/commit “2026_batch-quant-sum-stats_multi-way” (archived at https://github.com/SCohenLab/Extended-analysis-for-infer-subc-v2.0.0b1). The pipeline continues to be actively developed, and the updated implementation is available at (https://github.com/SCohenLab/infer-subc/blob/v2.0.0b1/notebooks/part_2_quantification). Per-object measurements were quantified from each organelle, or combination of organelles (contact sites). These metrics were then summarized on a per-subcompartment basis, yielding a total of 5,437 organelle metrics for soma regions and 3,316 metrics for neurites. The lower number of metrics included in the neurite analysis is due to the exclusion of organelle and contact site distribution analysis.

To reduce redundancy and improve statistical power, soma metrics were narrowed to a curated set of 248 features, neurite metrics to 117 features, and for the comparison of soma and neurite to 114 features (first pass metrics). These curated datasets were used for principal component analysis (PCA), hierarchical clustering and downstream statistical analyses (Data S1-3).

#### Organelle contact analysis

All possible organelle combinations of order k (k = 2 to N, where N is the number of segmented organelle classes) were enumerated, generating pairwise and higher-order sets (e.g., ER-mitochondria, ER-mitochondria-lysosome; known as “contact types”). For each contact type, 3D contact sites were computed as the overlap between included organelles, and each separate region of overlap was labeled to define the individual contact site. For each contact type, contact metrics were extracted, including contact site count (number of overlap components) and contact site volume.

To distinguish which contact sites were embedded within higher-order assemblies (e.g., ER-mitochondria contact sites that are within ER-mitochondria-peroxisome or other 3-way+ contact sites), each interaction site was screened for redundancy by testing whether the same overlap region intersected additional organelle segmentations not included in the current combination. Overlap components intersecting additional organelles were classified as embedded, whereas those not intersecting additional organelles were classified as exclusive.

To quantify contact order, analyses were restricted to exclusive contact sites only, thereby avoiding double-counting of shared overlap regions. Exclusive contacts were grouped by order (2-, 3-, 4-, and, when present, 5-way), and total contact number and contact volume were computed per subcompartment (soma or neurites) in each cell. Per-subcompartment summary statistics (mean, median, standard deviation) were calculated for each contact type. The pipeline used to quantify the order of the contacts, cell fraction are available at GitHub (“k-way_with_cell_fraction” https://github.com/SCohenLab/Extended-analysis-for-infer-subc-v2.0.0b1). The pipeline used to visualize the multi-way contact volumes is available at GitHub (“degree of interactions masks” https://github.com/SCohenLab/infer-subc/blob/v2.0.0b1/notebooks/part_2_quantification/method_interactions.ipynb).

#### Object-level contact, partner prevalence, and degree

Organelle contact at the object-level was analyzed using a complementary framework implemented in a custom Python/Jupyter pipeline. This analysis was based on the output of the infer_subc.quantification.batch.batch_process_quantification. Interaction-type lists were parsed to identify the focal organelle object (the organelle object being queried in the analysis; e.g., a lysosome) and its partner organelle classes (the organelle classes contacted by that focal object; e.g., mitochondria or lipid droplets), generating a per-cell list of focal-object-to-partner relationships. ER was excluded as a focal class due to its near-unity object count per cell but was retained as a partner. For each organelle type (focal organelle), we determined the list of partner organelles that had at least one contact site with the focal organelle. Interaction types were classified in reference to each organelle type (the focal organelle) and any other organelle that contacts the focal organelle (partners).

Contact vs. contact-free classification: For each cell and organelle type, focal objects were classified as contacting if they appeared at least once in the interaction list (≥1 partner), or as contact-free otherwise. For each cell, the percentage of contacting and contact-free objects was computed relative to the total number of objects of that organelle in the same cell. Per-cell percentages were then aggregated across cells and visualized as stacked proportions (stacked histograms of contacts vs not in contact), with summary statistics reported as mean ± SD.

Organelle partner prevalence (contact-only): Analyses of partner prevalence were restricted to only contacting objects. We calculated the percentage of contacting objects that touched each partner organelle at least once. Because a single object can contact multiple partners, partner prevalence categories are non-exclusive, and per-cell totals may exceed 100%. Per-cell partner prevalence profiles were aggregated across cells and displayed as stacked proportions.

Organelle contact degree (contact-only): For each contacting focal object, contact degree was defined as the number of distinct partner organelle types contacted at least once (degree 1 when contacting one partner class, up to the maximum possible, here 6). For each cell and focal organelle type, the degree distribution was computed as the percentage of contacting objects at each degree. These per-cell degree distributions were then aggregated across cells and visualized as stacked proportions.

Conceptual distinction between contact-site and object-level analyses: Together, these analyses distinguish contact-site architecture, captured by exclusive multi-way overlap regions, from object-level engagement, which quantifies how frequently individual organelles participate in contacts and with how many distinct partners. This dual framework avoids conflating discrete contact-site complexity with global organelle participation and enables complementary insights into organelle network organization. The pipeline used to quantify “Object-level contact, partner prevalence, and degree” are available at GitHub (“contact_analysis_suite_v2” https://github.com/SCohenLab/Extended-analysis-for-infer-subc-v2.0.0b1).

#### Principle component analysis

Principle component analysis (PCA) was carried out using the GraphPad Prism 10.2.3 Windows, GraphPad Software, Boston, Massachusetts USA, www.graphpad.com. The curated list of 248 organelle signature metrics for soma and 118 metrics for neurites, were first preprocessed by removing variables with a standard deviation of 0 and then standardized so that each variable had a mean of zero and a standard deviation of 1. Principle components (PCs) were selected using parallel analysis (1000 Monte Carlo simulations; selection based on eigenvalues greater than the 95% percentile in simulation; random seed auto-selected).

#### Hierarchical clustering

For hierarchical clustering analysis, first-pass metrics were preprocessed as described in the PCA section and used to build time-course trajectories for each metric by aggregating single-cell measurements within each time point (median). For each metric, median trajectories were z-scored across time points (per-metric standardization). To emphasize temporal changes rather than absolute levels, we computed the first discrete difference of each z-scored trajectory across consecutive time points (Δz between adjacent time points). Metrics were then hierarchically clustered based on Euclidean distance computed on these Δz vectors (SciPy, scipy.spatial.distance.pdist) using Ward linkage (scipy.cluster.hierarchy.linkage; dendrogram ordering via dendrogram). Cluster memberships were obtained by cutting the tree to a fixed maximum number of clusters (fcluster, criterion maxclust). Results were visualized as clustermaps in Python (Seaborn/Matplotlib), and summary tables (including raw medians, z-scores, Δz values, and cluster IDs) were exported to Excel. This code is available at GitHub (“timecourse_clustering_firstpass_soma_v1” https://github.com/SCohenLab/Extended-analysis-for-infer-subc-v2.0.0b1).

#### Dynamic time warping (DTW) analysis

To perform the correlation of tether expression with organelle contact dynamics, we implemented an automated analysis pipeline in RStudio based on dynamic time warping (DTW). The code takes three input files: (i) a pairing table specifying the tether protein name and the associated contact pair (and whether the contact metric is count or volume), (ii) a tether-expression table reporting protein abundance across differentiation time points (three biological replicates per time point [42], and (iii) contact-summary tables reporting, for each time point and contact pair, the per-cell contact count. Contact tables were filtered to retain pairwise contacts only (contact names containing exactly one “X”). For each time point, contact metrics were aggregated across cells (mean, median, SD, SEM), and tether expression was aggregated across replicates (mean, median, SD, SEM). For each tether-contact pair, time-series vectors were standardized by z-scoring across time points (z = (x − mean)/SD, computed within each pair across all available time points) to place expression and contact metrics on a common scale for visualization and comparison. DTW distances between the standardized tether-expression series and the standardized contact series were computed using the dtw package, allowing elastic alignment of the temporal trajectories. Statistical significance of DTW similarity was assessed by a permutation test, in which the time-point order of one series was randomly permuted B times (default B = 200; user-adjustable) to generate a null distribution of DTW distances. This code is available at GitHub (“Tethers_Contacts_DTW_v3” https://github.com/SCohenLab/Extended-analysis-for-infer-subc-v2.0.0b1).

#### Transparency

The following codes (“k-way_with_cell_fraction”, “contact_analysis_suite_v2”, “timecourse_clustering_firstpass_soma_v1”, “Tethers_Contacts_DTW_v3”) were generated with assistance from ChatGPT (OpenAI; 5.2 Thinking_2026) and subsequently reviewed, edited, and validated by the author (MCZ) across multiple independent datasets.

### Graphical abstract generation

All the graphical abstracts (Fig.1A; 4H; 5M; S7A; S12) were generated using BioRender (https://app.biorender.com/).

### Statistics

All experiments were performed with a minimum of three independent biological replicates unless otherwise specified. Biological replicates corresponded to independent cell cultures with matched passage number. The number of replicates and statistical tests are indicated in the figure legends. Unless otherwise stated, data were assumed to be approximately normally distributed; normality was not formally assessed. For multiple-group comparisons, one-way or two-way ANOVA was used, as indicated in the figure legend (p ≤0.05*, p ≤0.005**, p ≤0.0005***, p ≤0.0001****). For microscopy experiments, iPSC fields of view were selected from cells located in the center of colonies (avoiding colony edges), whereas iNeuron fields of view were selected at random. Unless otherwise stated, statistical analyses, plots, and heatmaps were generated using GraphPad Prism 10 (RRID:SCR_002798; GraphPad Software, Boston, MA, USA).

## Data availability

Multispectral imaging datasets (raw, unmixed, deconvolved, and segmented images) are deposited in the BioImage Archive (S-BIAD2983, https://www.ebi.ac.uk/biostudies/bioimages/studies). Additional data supporting the findings of this study are available from the corresponding author upon reasonable request.

## Acknowledgments

We thank Dr. Bill Skarnes for generously providing the KOLF2.1J wild type cell line and for helpful advice on clone selection. We thank Dr. Michael Ward for kindly sharing piggyBac vectors and Dr. Yue Andy Qi for sharing proteomics data collected from iNeurons throughout differentiation. We thank Wendy Salmon for expert microscopy advice. We acknowledge the UNC Hooker Imaging Core Facility supported in part by P30 CA016086 Cancer Center Core Support Grant to the UNC Lineberger Comprehensive Cancer Center and the UNC Department of Chemistry Mass Spectrometry Core Laboratory supported by the National Science Foundation under Grant CHE-1726291, which supported lipidomic analyses. We thank members of the Cohen and Deshmukh labs for helpful discussions.

## Author contributions

Conceptualization: MCZ, MD, SC

Methodology: MCZ, ZC, SNR, NRM

Investigation: MCZ, DB, CH, RB, BE

Visualization: MCZ, DB

Funding acquisition: MD, SC

Project administration: MD, SC

Supervision: MD, SC

Writing - original draft: MCZ

Writing - review & editing: MD, SC

---

**Fig. S1.**
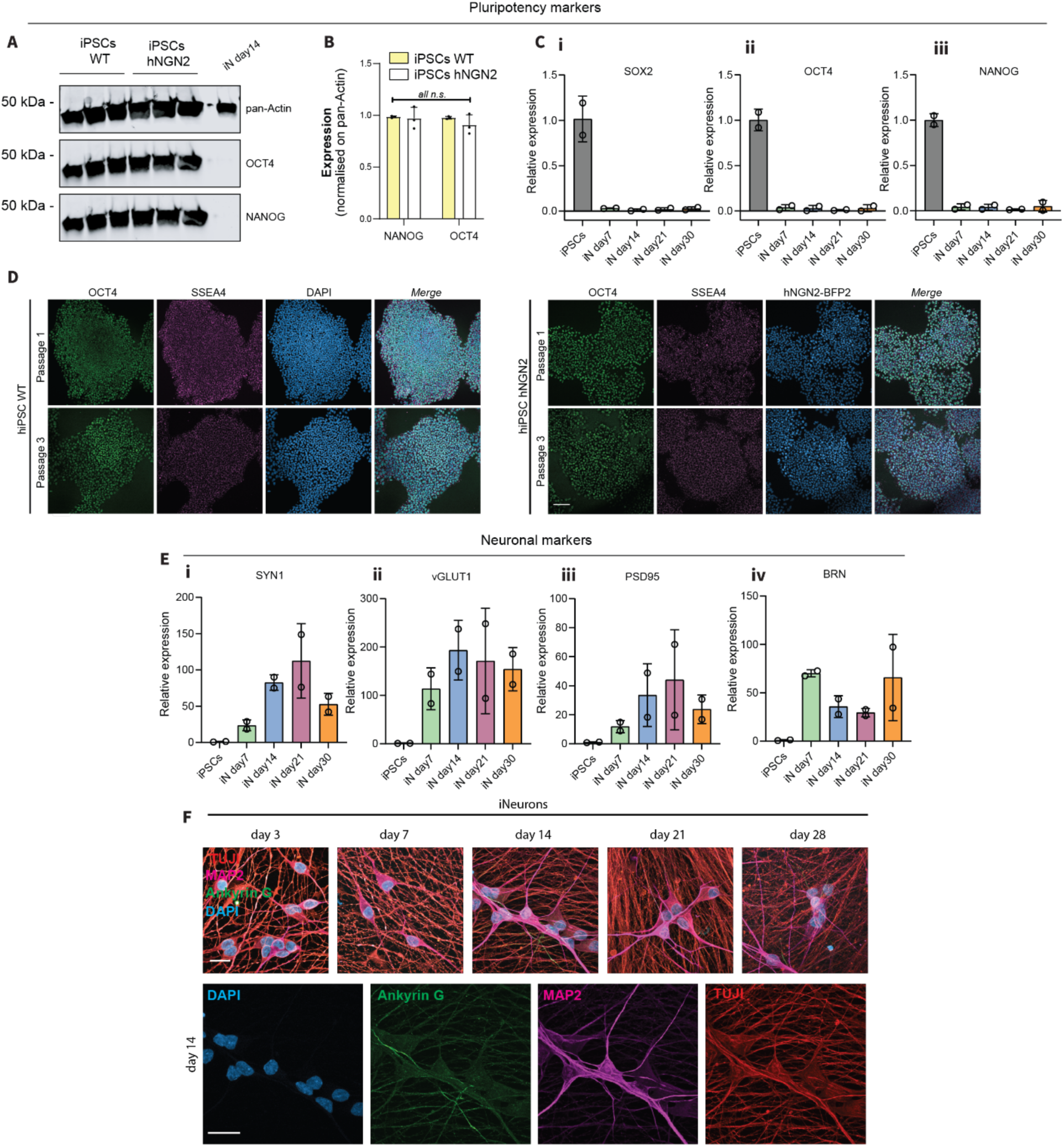
Clone generation and differentiation. (**A, B**) Immunoblot and its quantification, against pluripotent markers (OCT4, NANOG) and the loading control (pan-Actin) of wild-type hiPSCs (hiPSCs WT), hiPSCs integrated with hNGN2 (hiPSCs hNGN2), and day 14 iNeurons (differentiated by the same clone). (**C**) Real time qPCR of pluripotent genes as SOX2 (**Ci**), OCT4 (**Cii**), and NANOG (**Ciii**). N=2. (**D**) Immunofluorescence of wild-type hiPSCs (hiPSCs WT), and hiPSCs integrated with hNGN2 (hiPSCs hNGN2), against pluripotent markers (SSEA4, OCT4) and nuclei (DAPI). Scale: 100 µm. (**E**) Real time qPCR of neuronal genes as SYN1 (**Ei**), VGLUT1 (**Eii**), PSD95 (**Eiii**), and BRN (**Eiv**). (**F**) Immunofluorescence of neuronal markers (Ankyrin G; MAP2; and Beta-III-Tubulin (TUJI)), during the differentiation into forebrain-like neurons at different time points. Scale: 10 µm.

**Fig. S2.**
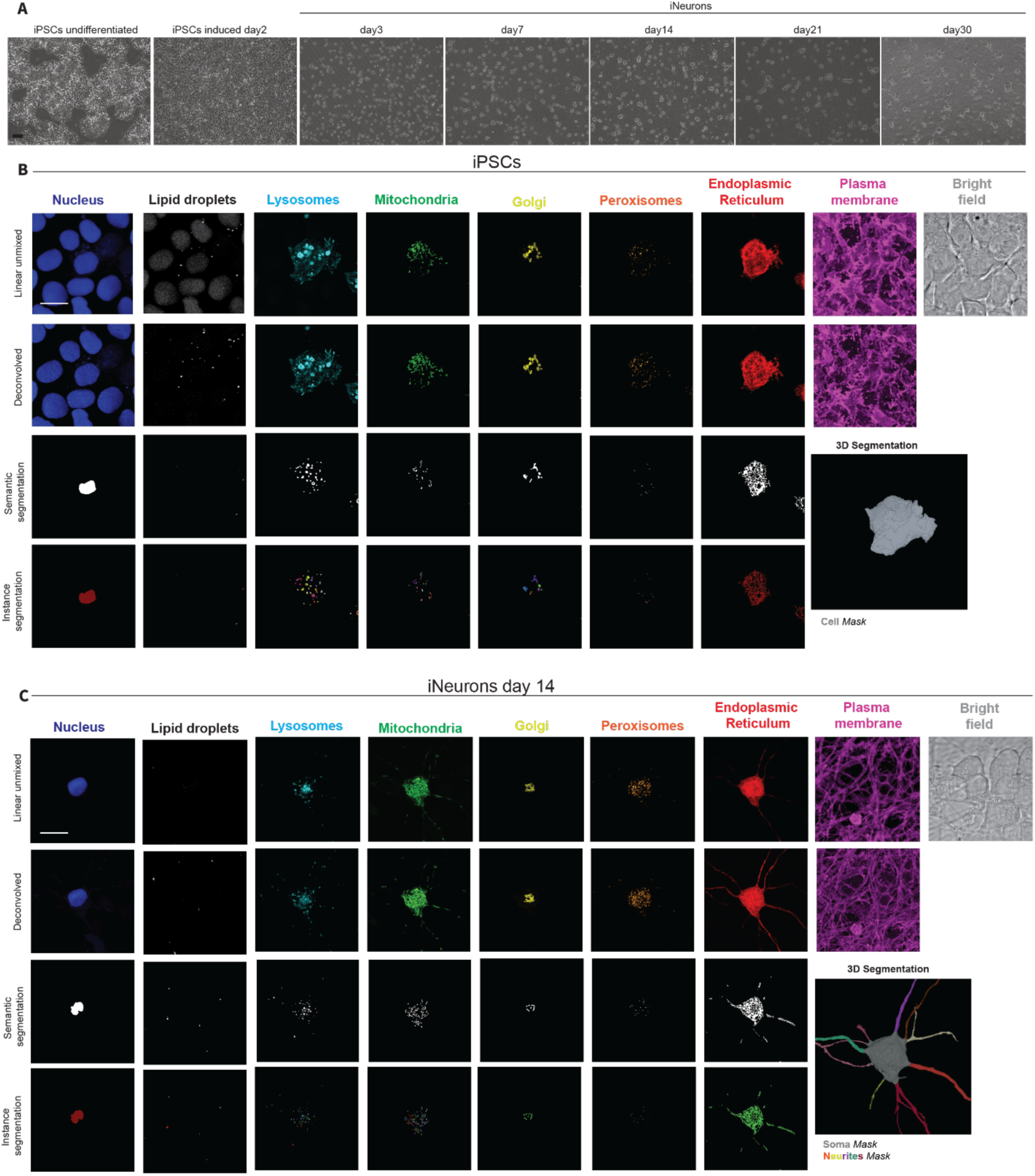
Multispectral imaging and segmentation. (**A**) Bright field images of iPSCs and iNeurons along differentiation. Scale: 100 µm. (**B**) Multispectral imaging of iPSCs. (**C**) Multispectral imaging of iNeurons. B-C include linear unmixed and deconvolved images (maximum intensity projections), as well as semantic and instance segmentations of organelles (single z plane), and 3D masks of the cell (**B**) or soma and neurites (**C**). Scale: 10 µm.

**Fig. S3.**
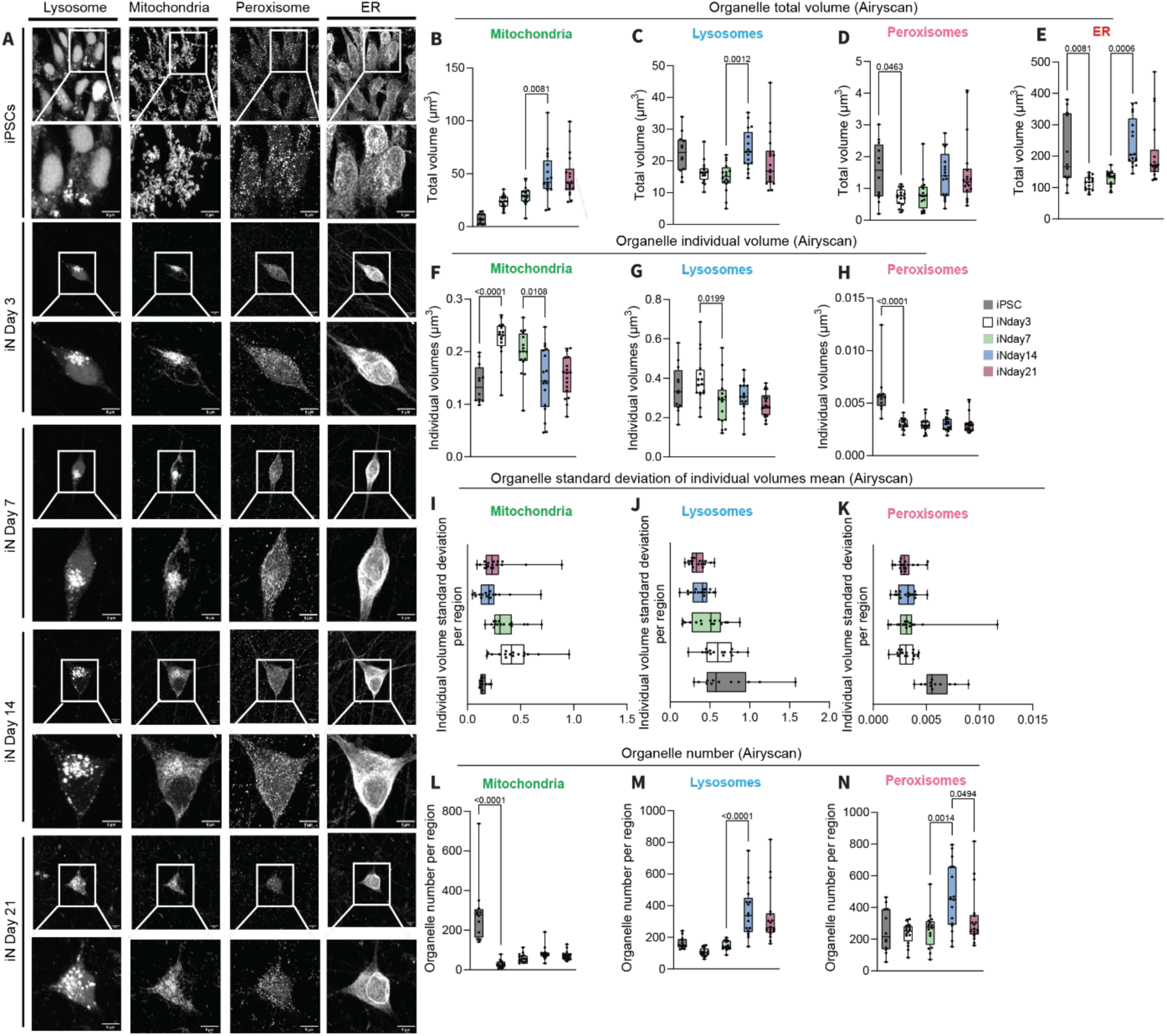
Airyscan of lysosomes, mitochondria, peroxisomes and ER. **(A)** Airyscan images of four organelles in fixed samples during neuronal differentiation (lysosomes, mitochondria, peroxisomes, and endoplasmic reticulum (ER)). Scale: 5 µm. (**B-E**) Total volume of mitochondria, lysosomes, peroxisomes and ER, per cell (iPSCs) or soma (iNeurons) region, in iPSCs and iNeurons at day3, day7, day14 and day21. (**F, G, H**) Individual organelle volumes of mitochondria, lysosomes, peroxisomes, and their associated standard deviation (**I, J, K**), and number (**L, M, N**) per region, in iPSCs and iNeurons at day3, day7, day14 and day21. One-way ANOVA, p-values versus previous time point.

**Fig. S4.**
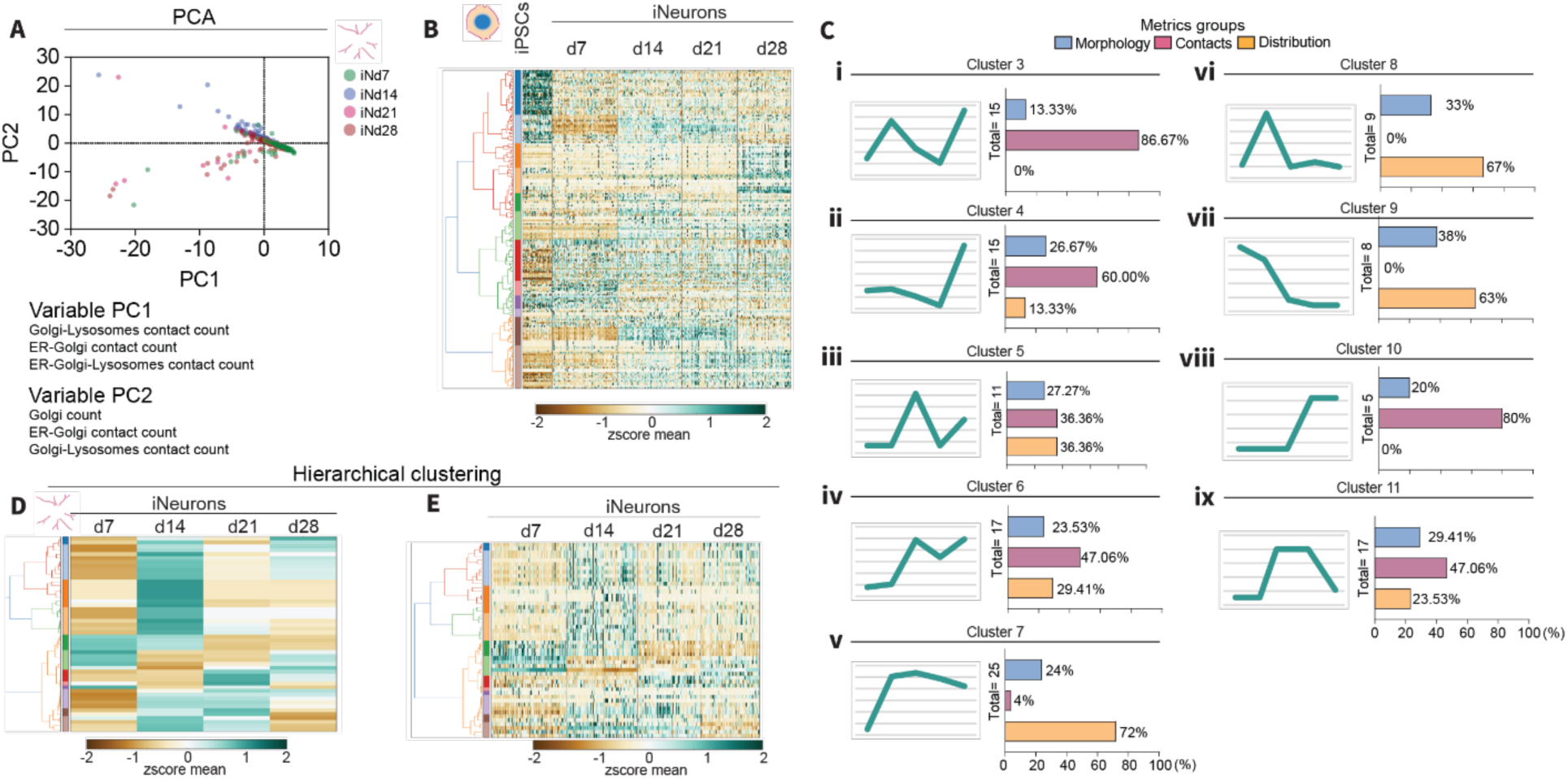
PCA and hierarchical clustering of neurites. (**A**) PCA of 120 metrics of the neurite region, with the top three variables in PC1 and PC2. (**B**) Hierarchical clustering of 248 metrics of the soma region during neuronal differentiation at single cell resolution. (**C**) Curves representing the dynamics of a representative metric and graphs showing percentage of each metric group (morphology, contacts and distribution of the organelle) within each cluster. (**D, E**) Hierarchical clustering of 117 metrics of the neurite region as a median er time point (**D**) or at single cell resolution (**E**).

**Fig. S5.**
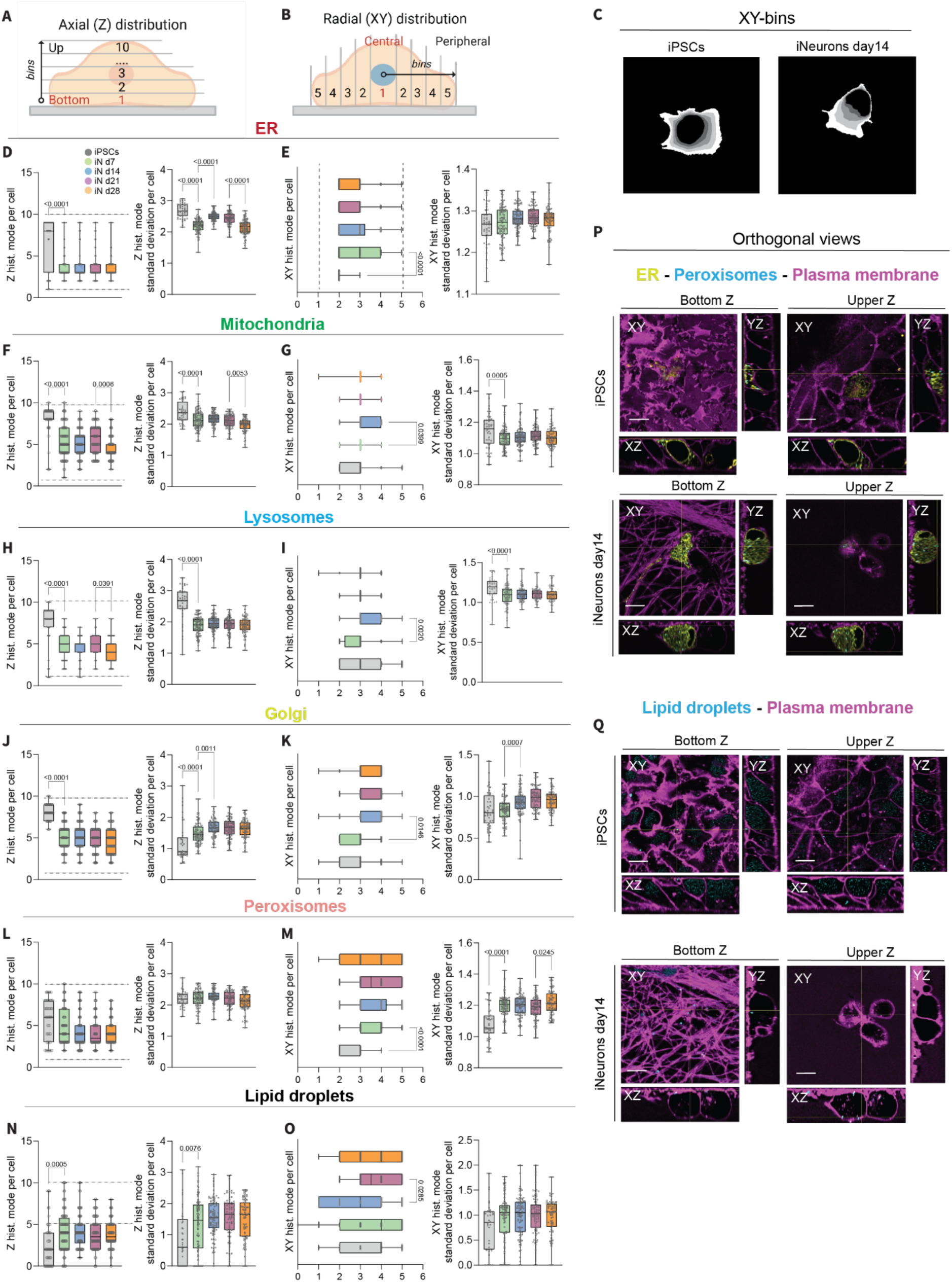
Distribution of the organelles within the soma region, during neuronal differentiation. (**A**) Graphical representation of the axial (Z) and (**B**) lateral (XY) distributions of the organelles in the cell or soma region. (**C**) XY-bin images of an iPSC and the soma region of an iNeuron at day14. (**D, E**) Axial and lateral distribution, with standard deviation, per region, of the ER; (**F, G**) mitochondria; (**H, I**) lysosomes; (**J, K**) Golgi; (**L, M**) peroxisomes; (**N, O**) lipid droplets. (**P, Q**) Deconvolved multispectral images (maximum intensity projection) of iPSCs and iNeurons at day14, showing orthogonal projections (in XY, XZ and ZY) of ER (yellow), peroxisomes (cyan) and plasma membrane (magenta); (**Q**) and lipid droplets (cyan), together with plasma membrane (magenta). Scale: 10 µm. One-way ANOVA, p-values versus previous time point.

**Fig. S6.**
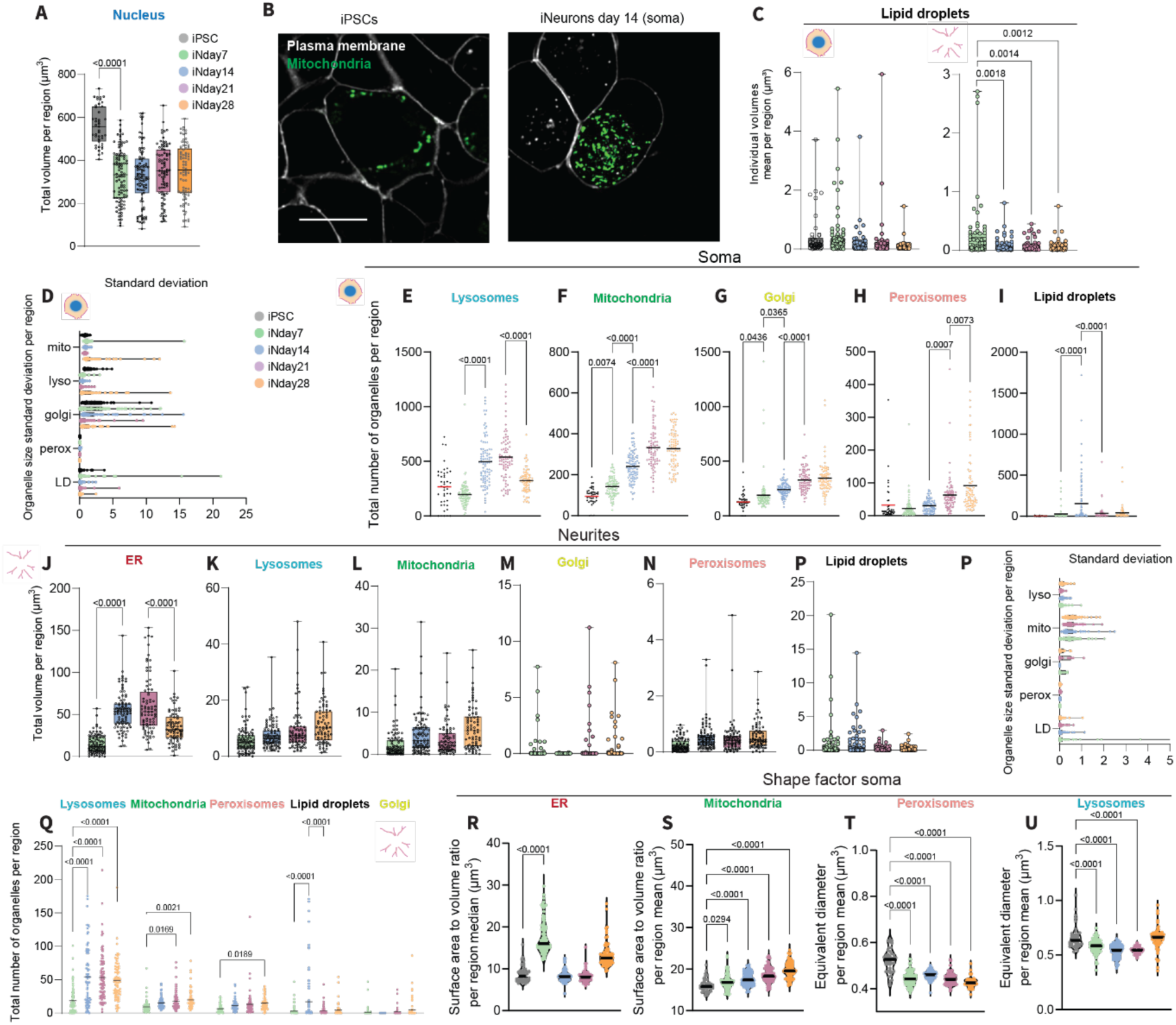
Extended organelle profiling in iNeuron soma and neurite regions. (**A**) Total volume of the nucleus during neuronal differentiation. (**B**) Deconvolved multispectral images (single z plane) of iPSCs and iNeurons at day14, showing plasma membrane channels (grey), together with mitochondria (green). Scale: 10 µm. (**C**) Mean volume of individual lipid droplets, in the soma and neurite regions, during neuronal differentiation. (**D**) Individual organelle volume mean in the soma region during neuronal differentiation. (**E-I**) Total number of lysosomes, mitochondria, Golgi, peroxisomes and lipid droplets per soma region during neuronal differentiation. (**L-O**) Total volume of ER, lysosomes, mitochondria, Golgi, peroxisomes and lipid droplets per cell in the neurite regions during neuronal differentiation. (**P**) Individual organelle volume mean standard deviation in the neurite regions during neuronal differentiation. (**Q**) Total number of lysosomes, mitochondria, Golgi, peroxisomes and lipid droplets per cell in the neurite region during neuronal differentiation. Two-way ANOVA. (**R-U**) Shape factor of ER, mitochondria, peroxisomes, and lysosomes in the soma region during neuronal differentiation. One-way ANOVA, p-values versus previous time point.

**Fig. S7.**
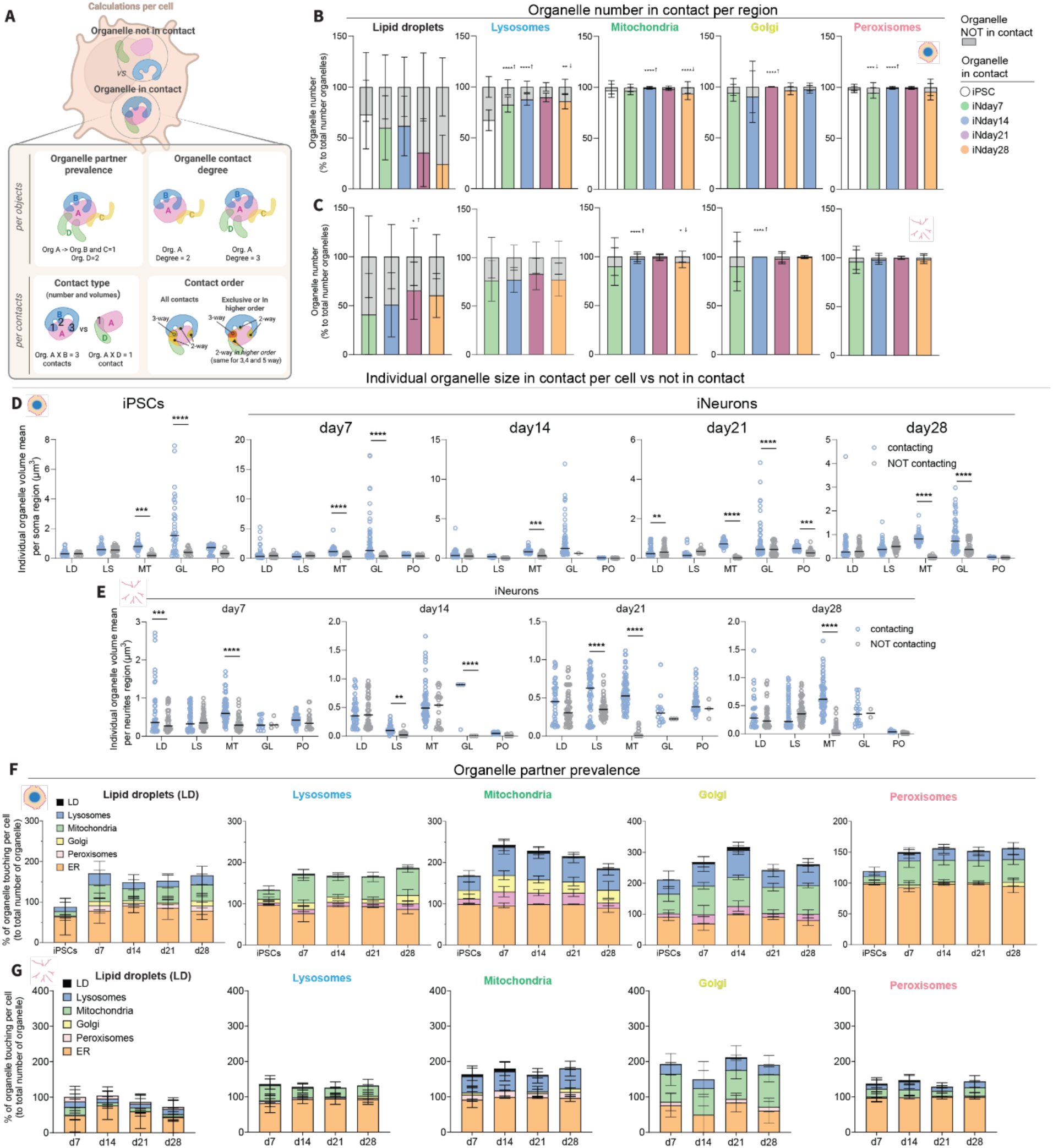
Organelle object contact profiling and extended contact order analysis. (**A**) Illustration of the contact analysis per cell. Each measurement is shown as per region measurements (whole cell for iPSCs and soma or neurites for iNeurons), per time point. For each organelle we measured the number of objects that are in contact, or not, with at least one other organelle (“Contacting organelles” versus “NOT Contacting organelles”). From these objects we measured their individual size and plotted the mean per cell or soma region (“Individual organelle size”). From each organelle object, we measured how often each organelle type contacted each of the others at least once per region (“Organelle partner prevalence”); and the number of distinct organelle types that a given organelle object contacts at least once, from degree 1 (one type) up to degree 5 (five types) (“Organelle contact degree”). For each contact, we first defined its type (e.g., ER-lysosomes, ER-mitochondria) and then quantified the number of overlapping surfaces and their volumes (“Contact type”). We then defined each contact’s “Contact order” by assessing whether it formed an exclusive 2-, 3-, 4-, or more rarely, 5-way contact, excluding contacts embedded within higher-order assemblies. (**B**) Percentage of the organelle objects in contact versus not, in the soma or (**C**) neurite regions. (**D**) Individual organelle mean volume in contact versus not, in the soma or (**E**) neurite regions. (**F**) Organelle partner prevalence, in percentage to total number of contacts per cell, in the soma region. (**G**) Organelle partner prevalence of the neurite region. Two-way ANOVA, p-value versus previous time point.

**Fig. S8.**
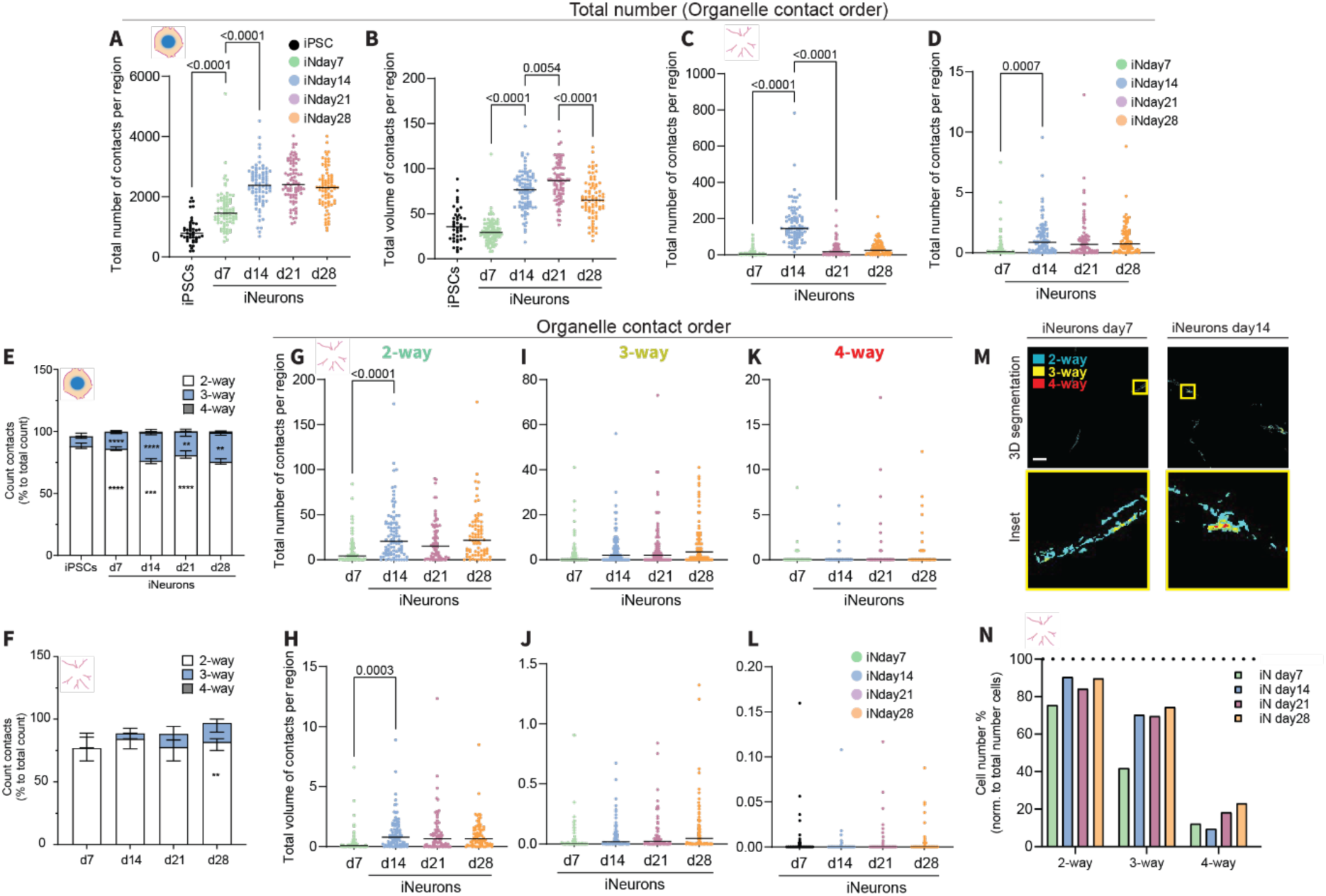
Extended organelle contact order profiling along differentiation. (**A, B**) Total number of contacts (exclusive) or (**C, D**) volume of contacts, per cell in the soma or neurite regions. (**E**) Contact order fraction, per cell, along neuronal differentiation, in the soma and (**F**) neurite region. (**G-L**) Organelle contact order (2-, 3-, 4-way) number and volumes of the neurite region. (**M**) Imaging of 2-, 3-, and 4-way contact volumes, in the neurite region of iNeurons at day 7 and 14. Scale: 10 µm. (**N**) Number of cells, in percentage, per time point, that has at least one contact 2-, 3-, or 4-way. Two-way ANOVA, p-values versus previous time point.

**Fig. S9.**
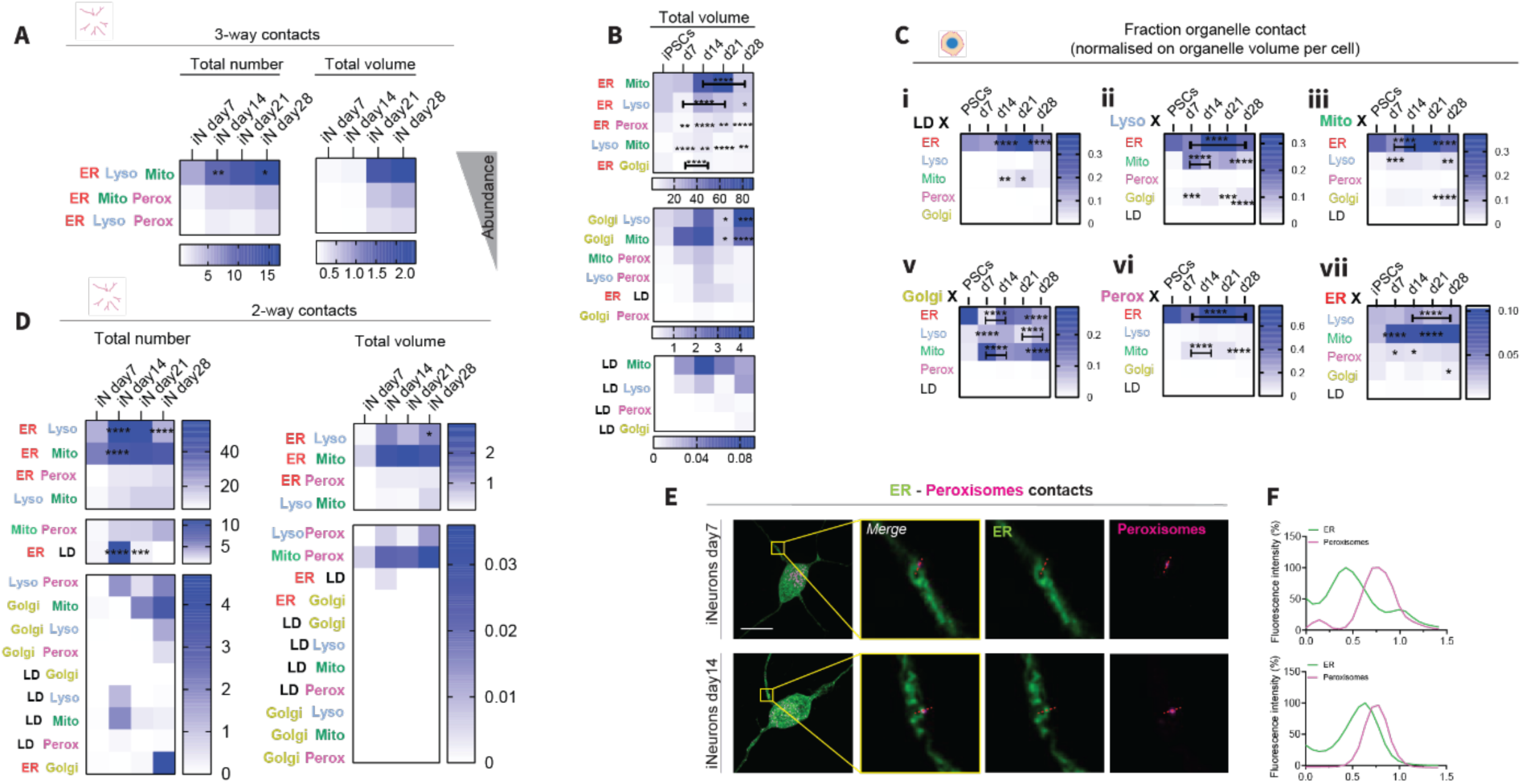
Extended organelle contact type profiling along differentiation. (**A**) Total number and volume of 3-way contacts, per cell in the neurite region. (**B**) Organelle contact pair total volumes (not normalized), per cell in the soma region. (**C**) Organelle contact pair total volume fraction, per cell in the soma region. (**D**) Total number and volume of the 2-way contacts, per cell in the neurite region. (**E**) Multispectral imaging and (**F**) line scans of ER-peroxisome contacts in the neurite region of iNeurons at day 7 and day 14. Scale: 15 µm. Two-way ANOVA, p-values versus previous time point.

**Fig. S10.**
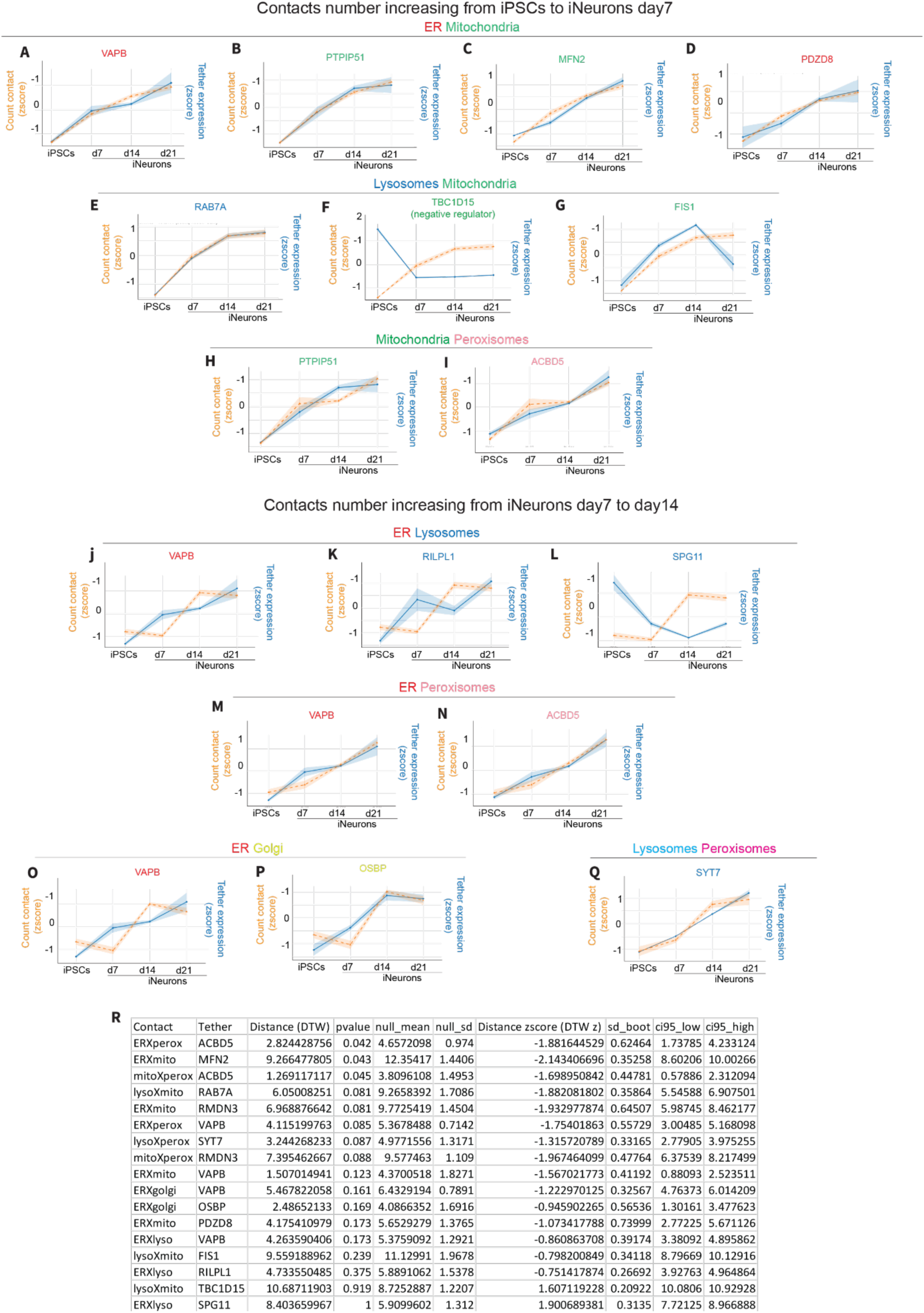
The number of contacts per cell correlate with the expression of known protein tethers. Dynamic time warping (DTW) measuring the correlation between the number of contact pairs and their associated tether proteins. (**A-D**) Correlation of lysosome-mitochondria contact number with VAPB, PTPIP51, MFN2 or PDZD8. (**E-G**) Correlation of ER-mitochondria contact number with RAB7A, TBCD15, or FIS1. (**H-I**) Correlation of mitochondria-peroxisome contact number with PTPIP51 or ACBD5. (**J-L**) Correlation of ER-lysosome contact number with VAPB, RILPL1 or SPG11. (**M-N**) Correlation of ER-peroxisome contact number with VAPB or ACBD5. (**O-P**) Correlation of ER-Golgi contact number with VAPB or OSBP. (**Q**) Correlation of lysosome-peroxisome contact number with SYT7. Tether expression data from reference [42].

**Fig. S11.**
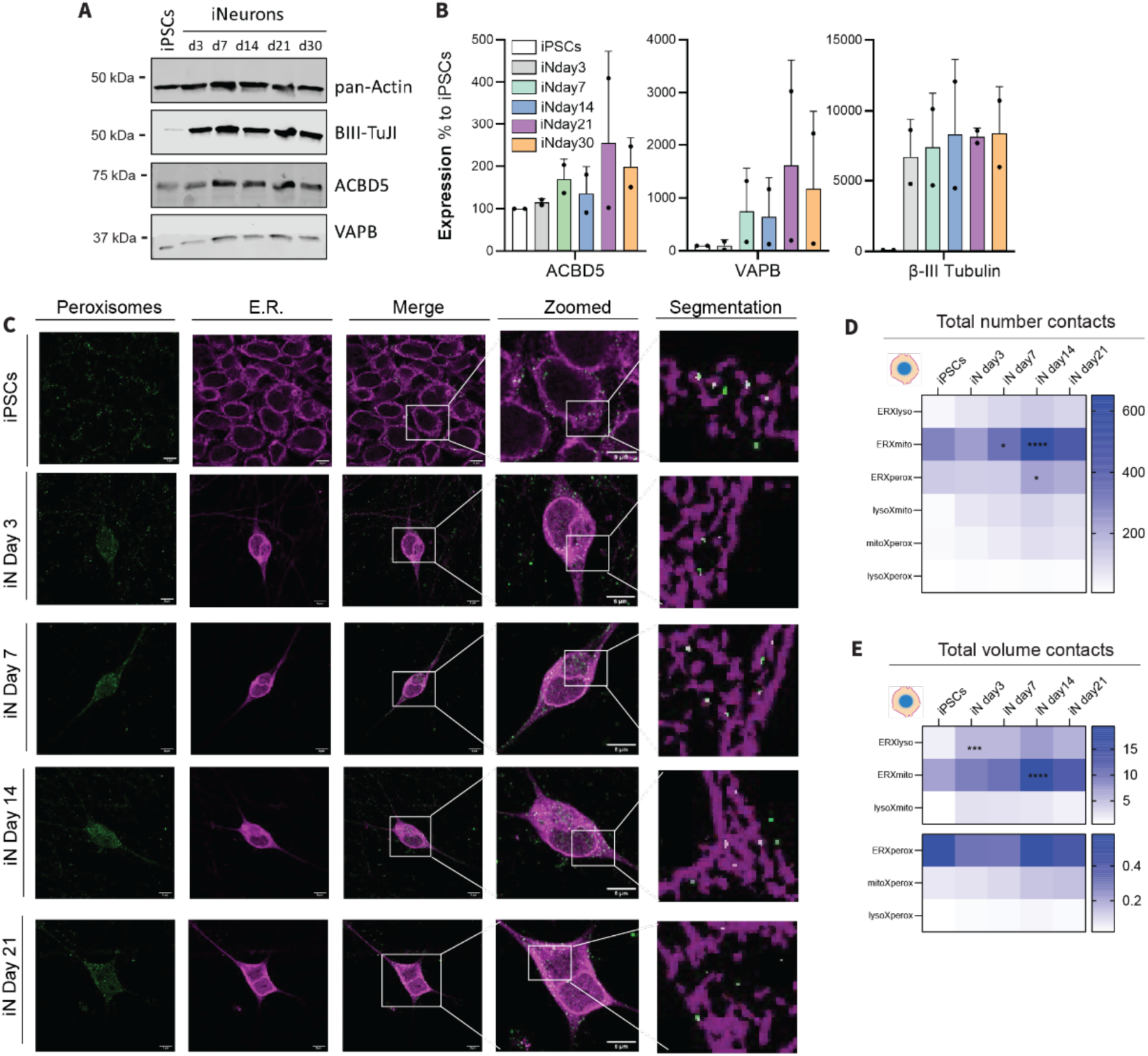
Validation of ER-peroxisome tethers and contacts. (**A**) Immunoblot, and its (**B**) quantification, of the tethering proteins VAPB and ACBD5 with loading control (pan-actin) together with a neuronal marker (BIII-TuJI). N= 2. (**C**) Immunofluorescence AiryScan images of endogenously stained peroxisomes and ER. (**D**) Total number and (**E**) volume of all the contact pairs per soma region along neuronal differentiation by AiryScan imaging. Scale: 5 µm. Two-way ANOVA, p-values versus previous time point.

**Fig. S12.**
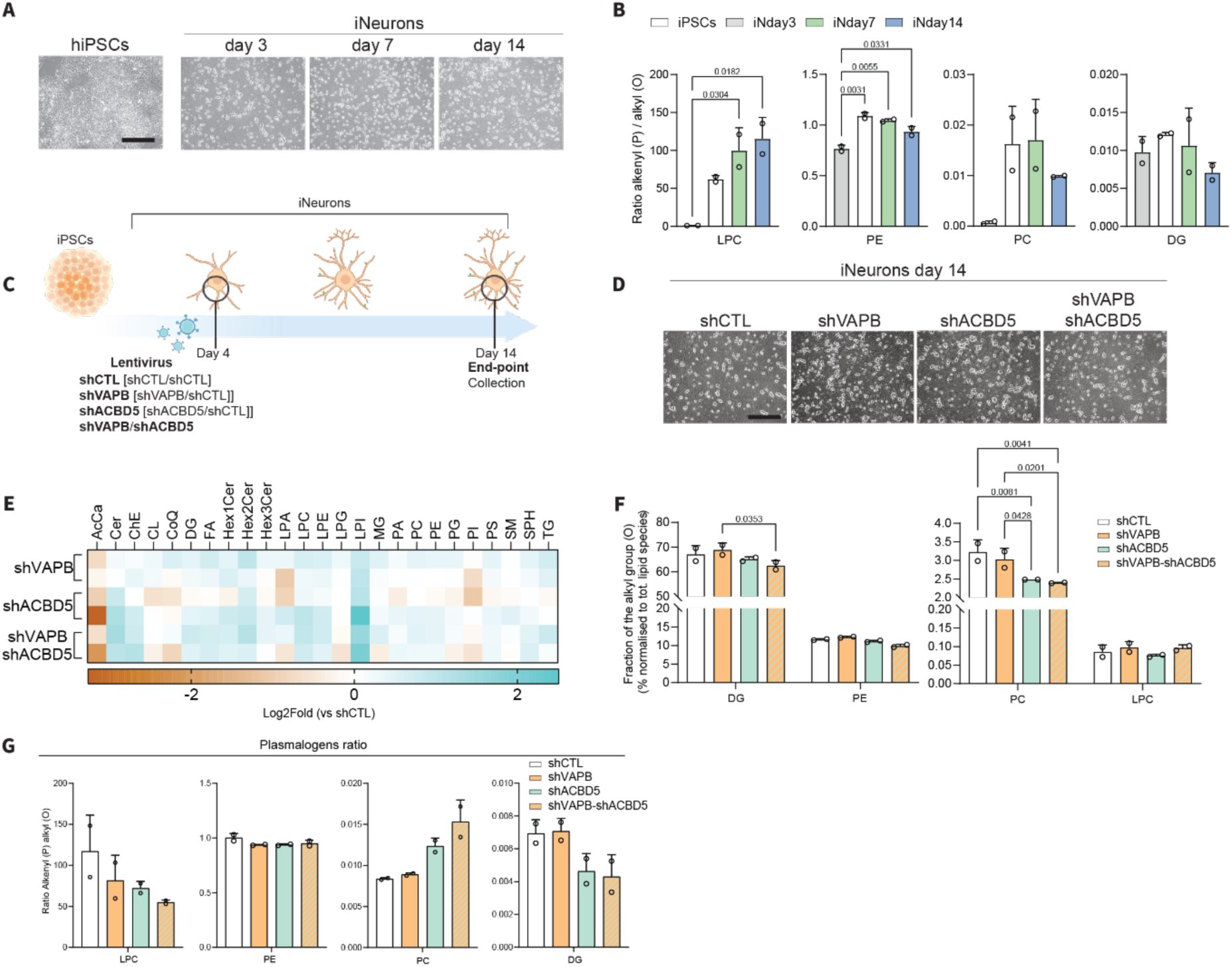
Lipidomic analysis. (**A**) Brightfield images of the cells along neuronal differentiation before collecting for lipidomic analysis.(**B**) Ratio of the alkenyl (P) group with the alkyl (O) group, at different time points during neuronal differentiation (iPSCs, iNeurons at day3, day7 and day 14), N=2. One-way ANOVA. (**C**) Graphical workflow of the knock-down of VAPB and ACBD5, alone or in combination, in iNeurons. (**D**) Brightfield images of the cells upon knock-down of VAPB and ACBD5, alone or in combination, at day 14 of differentiation, before collecting for lipidomic analysis. Scale: 300 µm (**E**) Total lipid amount in iNeurons at day 14 after knock-down of VAPB and ACBD5, alone or in combination, normalized to the scramble control (Log2 fold). (**F**) Fraction of the alkyl (O) group upon knock-down of VAPB and ACBD5, alone or in combination, at day 14 of differentiation. (**G**) Ratio of the alkenyl (P) group with the alkyl (O) group, upon knocking-down VAPB and ACBD5, alone or in combination, at day 14 of differentiation. N=2. Two-way ANOVA.

**Fig. S13.**
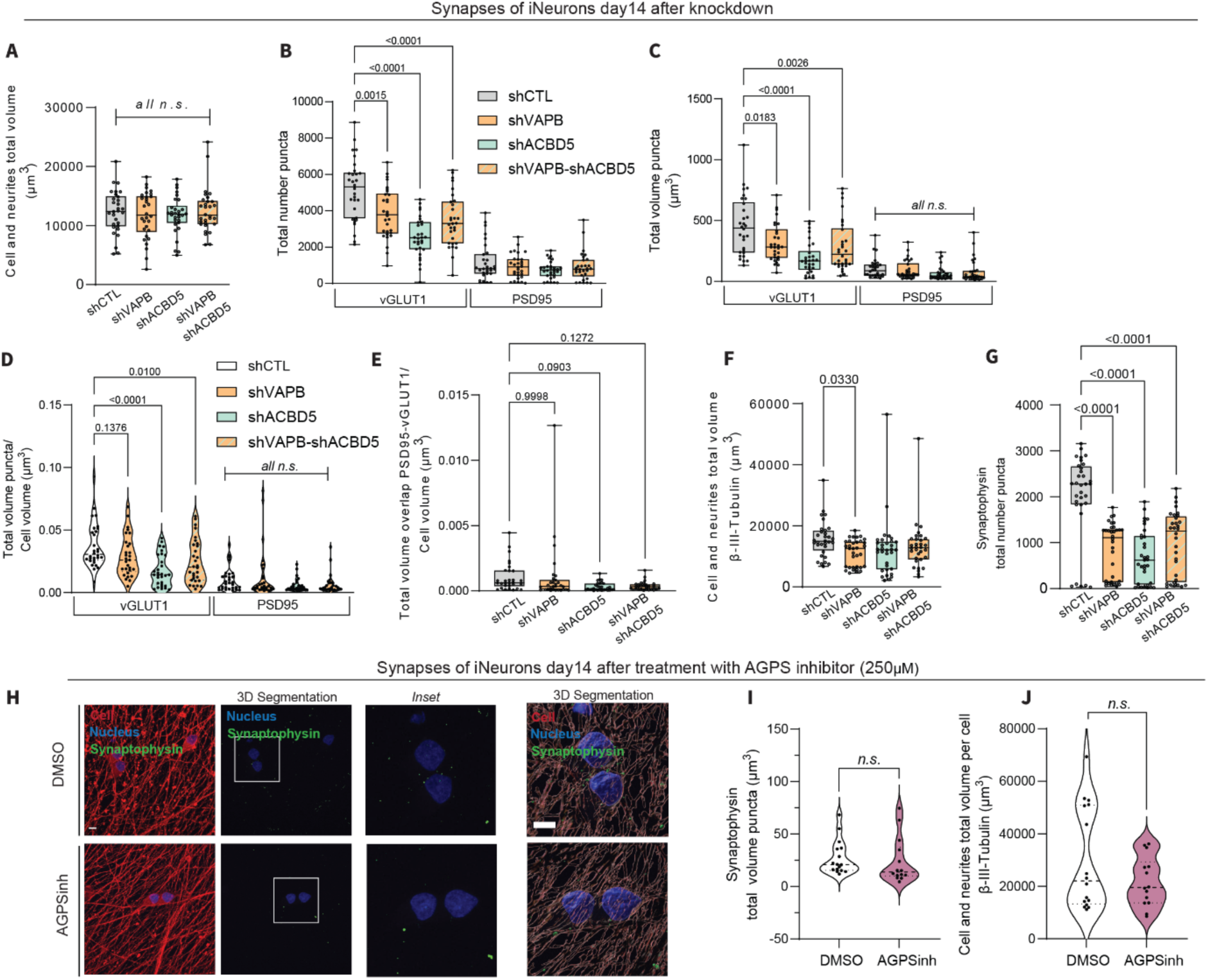
Extended synapse analysis. (**A**) Cell and neurite volumes measured as total volume from MAP2 signal, of iNeurons at day 14, after knock-down of VAPB and ACBD5 alone or in combination, together with the scramble Control. (**B-E**) Synapses quantification by measuring the number, volume, and overlap of PSD95 and vGLUT1 puncta, of iNeurons at day 14, after knock-down of VAPB and ACBD5 alone or in combination, together with the scramble Control. (**F**) Cell and neurite volumes measured as total volume from β-III-Tubulin signal, of iNeurons at day 14, after knock-down of VAPB and ACBD5 alone or in combination, together with the scramble Control. (**G**) Synapses quantification by measuring the number of Synaptophysin puncta, in iNeurons at day 14, after knock-down of VAPB and ACBD5 alone or in combination, together with the scramble Control. (**H, I**) Synapses quantification by measuring the number of Synaptophysin puncta, in iNeurons at day 14 treated with AGPS inhibitor. (**J**) Cell and neurite volumes measured as total volume from β-III-Tubulin signal, in iNeurons at day 14 treated with AGPS inhibitor. Unpaired t-Test. Scale: 10 µm.

**Data S1.**

All first pass metrics of iPSCs and the soma region of iNeurons at day 7, 14, 21 and 28. This file contains the tabulated data behind the figure 1D.

**Data S2.**

All first pass metrics of the soma and neurites region metrics of iNeurons at day 7, 14, 21 and 28. This file contains the tabulated data behind the figure 1E.

**Data S3.**

All first pass metrics of the neurites region metrics of iNeurons at day 7, 14, 21 and 28. This file contains the tabulated data behind the figure S4A.

**Data S4.**

Hierarchical clustering of iPSCs and soma region metrics of iNeurons at day 7, 14, 21 and 28. This file contains the tabulated data behind the figure 1F.

**Data S5.**

Hierarchical clustering of the neurite’s region metrics of iNeurons at day 7, 14, 21 and 28. This file contains the tabulated data behind the figures S4D, E.

**Data S6.**

Organelle object extended profiling. This file contains the tabulated data behind the figure S7, and figure 3A.

**Data S7.**

Organelle contact order extended profiling. This file contains the tabulated data behind the figures 3B to G, I and figures S8.

**Data S8.**

Lipidomics during neuronal differentiation, and at iNeurons day14, after knockdown of VAPB and ACBD5. This file contains the tabulated data behind the figures 4I, 5C to E and figures S12.

